# Measuring Directed Functional Connectivity Using Non-Parametric Directionality Analysis: Validation and Comparison with Non-Parametric Granger Causality

**DOI:** 10.1101/526566

**Authors:** Timothy O. West, David M. Halliday, Steven L. Bressler, Simon F. Farmer, Vladimir Litvak

## Abstract

**Background:** ‘Non-parametric directionality’ (NPD) is a novel method for estimation of directed functional connectivity (dFC) in neural data. The method has previously been verified in its ability to recover causal interactions in simulated spiking networks in Halliday et al. (2015)

**Methods:** This work presents a validation of NPD in continuous neural recordings (e.g. local field potentials). Specifically, we use autoregressive model to simulate time delayed correlations between neural signals. We then test for the accurate recovery of networks in the face of several confounds typically encountered in empirical data. We examine the effects of NPD under varying: a) signal-to-noise ratios, b) asymmetries in signal strength, c) instantaneous mixing, d) common drive, e) and parallel/convergent signal routing. We also apply NPD to data from a patient who underwent simultaneous magnetoencephalography and deep brain recording.

**Results:** We demonstrate that NPD can accurately recover directed functional connectivity from simulations with known patterns of connectivity. The performance of the NPD metric is compared with non-parametric Granger causality (NPG), a well-established methodology for model free estimation of dFC. A series of simulations investigating synthetically imposed confounds demonstrate that NPD provides estimates of connectivity that are equivalent to NPG. However, we provide evidence that: i) NPD is less sensitive than NPG to degradation by noise; ii) NPD is more robust to the generation of false positive identification of connectivity resulting from SNR asymmetries; iii) NPD is more robust to corruption via moderate degrees of instantaneous signal mixing.

**Conclusions:** The results in this paper highlight that to be practically applied to neural data, connectivity metrics should not only be accurate in their recovery of causal networks but also resistant to the confounding effects often encountered in experimental recordings of multimodal data. Taken together, these findings position NPD at the state-of-the-art with respect to the estimation of directed functional connectivity in neuroimaging.

**Highlights:**

- Non-parametric directionality (NPD) is a novel directed connectivity metric.
- NPD estimates are equivalent to Granger causality but more robust to signal confounds.
- Multivariate extensions of NPD can correctly identify signal routing.

**Abbreviations:**

dFC: Directed functional connectivity
EEG: Electroencephalogram
LFP: Local field potential
MEG: Magnetoencephalogram
MVAR: Multivariate autoregressive (model)
NPD: Non-parametric directionality
NPG: Non-parametric Granger (causality)
SMA: Supplementary motor area
SNR: Signal-to-noise ratio
STN: Subthalamic Nucleus

## 1 Introduction

Questions regarding the causal relationships between anatomically connected regions of the brain have become of fundamental importance across many domains of neuroscience (Sporns 2010; Swanson 2012). A novel method for estimating directed functional connectivity (dFC), termed non-parametric directionality (NPD), has been recently described in Halliday (2015). This method has been demonstrated to yield physiological insights into the connectivity of the cortico-basal-ganglia network when applied to (continuous) field recordings made in rodents (West et al. 2018). In this work we evaluate NPD’s performance at recovering known patterns of connectivity in the face of several confounding factors and compare it with another popularly used metric – Granger causality.

Functional connectivity is based on a description of the statistical dependencies between different neural signals and is typically estimated through time or frequency domain correlations (Friston 2011; Bastos and Schoffelen 2016). Magnitude squared coherence, equivalent to a frequency domain coefficient of correlation, has been widely adopted as the estimator of choice for functional connectivity in the neuroimaging community (Brillinger 1975; Halliday et al. 1995). Undirected measures of functional connectivity (such as coherence) are symmetrical, giving no indication of the temporal precedence of correlations, a property understood to underlie causation in time evolving systems (Wiener 1956), nor the predictability of one time series from that of the other. dFC aims to estimate statistical asymmetries in the correlated activity of a set of signals in order to infer the causal influence (or predictability) of one signal over another. Similar to the role played by coherence in measuring undirected functional connectivity, Wiener-Granger causality has emerged as a first-choice estimator of directed connectivity due to its well established theoretical basis (Ding et al. 2006; Bressler and Seth 2011) and its successful application to questions concerning causal networks inferred from large-scale neural recordings (e.g. Brovelli et al. 2004; Richter et al. 2018).

Estimates of dFC are most frequently computed in the literature using Granger causality or one of its variants (Granger 1969; Geweke 1982; Kamiński et al. 2001; Dhamala et al. 2008). Granger causality is expressed in terms of the capacity of the information in one signal’s past to predict the future of another signal. Granger (1969) introduced a straightforward measurement method through the implementation of an autoregressive model by which the explained variance of *Y* is compared between that of a ‘full’ model (i.e. accounting for the past of *X* and *Y*) and that of a restricted model (i.e. *Y* only). If a prediction of the future of *Y* is aided by information from the past of *X*, then *X* is said to ‘Granger-cause’ *Y*. The method requires factoring out the autoregressive component of the signal (i.e. the ‘restricted’ model) to avoid trivial correlations that occur simply due to the periodicity in the signals.

Efforts to estimate Granger causality without resorting to autoregressive models have resulted in an extension of Granger causality termed non-parametric Granger causality (NPG), which avoids the estimation of transfer functions from multivariate autoregressive (MVAR) coefficients (Dhamala et al. 2008). In NPG, transfer functions and noise covariances are estimated through the spectral factorization of (non-parametrically derived) Fourier coefficients rather from MVAR model parameters. Here, we directly compare NPG with NPD as an estimator of dFC. Both methods share the property of being non-parametric (model-free) approaches which can be derived from identical spectral transforms made either via Fourier or wavelet techniques.

NPD is founded on the same principles of causality as Granger, namely that temporally lagged *dependencies* indicate causal direction. NPD works by decomposing the coherence into three temporally independent components separated by the relative lag of the dependencies between the signals: 1) forward lagged; 2) reverse lagged; and 3) instantaneously correlated. Rather than using a naïve cross-correlation estimator that is susceptible to spurious peaks due to the individual signals’ autocorrelations, NPD takes an approach akin to the factoring out of a ‘restricted’ model (*Y* only) used in Granger. This is achieved through a process of spectral pre-whitening which acts to bring the individual signals spectra closer to white-noise but preserves the correlations between them. In the original paper (Halliday 2015), the method was validated using a simple three node network with each node’s dynamics simulated using a conductance model of a spiking neurone in order to generate a series of discrete point processes. The authors demonstrated that NPD was successful in recovering the connectivity from a range of simulated architectures. Furthermore, the method was applied to spike timings (a point process) recorded from muscle spindle and shown to yield physiologically plausible causality results. Our recent work has extended the application of NPD to continuous local field potential (LFP) recordings made from an *in vivo* preparation of the cortico-basal ganglia system (West et al. 2018).

Estimation of empirical dFC in continuous neural recordings such as the LFP or magneto/electroencephalogram (M/EEG) is complicated by a number of factors. These include: low and possibly unequal signal-to-noise ratios (SNRs), instantaneous volume conduction, common drive, signal routing via parallel but disjoint paths, and the presence of cyclic paths within a network. All pose potential confounds for all the metrics described here. The failure of Granger causality in the presence of large amounts of measurement noise is a well-established shortcoming (Newbold 1978) which becomes particularly acute in noisy electrophysiological recordings (Nalatore et al. 2007). Differences in the recording gain between signals is also known to confound estimation of Granger causality, with the metric being biased towards treating the strongest signal as the driver (Haufe et al. 2012; Bastos and Schoffelen 2016). This property is likely to be a nuisance when investigating causation between multimodal signal sets such as in experiments involving simultaneous measurements of MEG and LFP where significant differences in recording gain are to be expected (Litvak et al. 2011).

Instantaneous mixing of the electromagnetic signals generated by distinct sources in the brain has long been known to make estimation of functional connectivity based on recordings such as the EEG difficult (Nunez et al. 1997; Srinivasan et al. 2007; Haufe et al. 2012). Common presynaptic drive produces correlations in pairs of output spike trains (Farmer et al. 1993), and in pairs of evoked potentials (Truccolo et al. 2002). This problem can lead to spurious estimates of directed connectivity if delays in the arrival of the common input induce lagged correlations between unconnected neurons or neuronal populations. When the common presynaptic input is measured, extensions of functional connectivity metrics built upon partial regressions (so called *conditioned* or *partialized* estimates) can be used to remove common input effects and subsequently, remove the possibility of spurious inference of directed connectivity between neurones in receipt of lagged common input. Partial regression can be used with both NPD and NPG to reduce the influence of common drive. In the case of NPD, the authors introduced a multivariate extension that can be used to reduce the influence of common drive through partial regression of a third reference signal (Halliday et al. 2016). This method relies upon the reference signal substantially encapsulating the activity of the common drive. In the case that the recordings are incomplete representations of the propagating neural activity, the conditioning will only be partially effective. NPD and NPG conditioned on a third signal can also be used to infer connectivity patterns where two signals are correlated through interaction with an intermediary signal (West et al. 2018).

In this paper we will assess the performance of NPD’s ability to recover the connectomes from several simulated architectures and in the face of the previously stated confounds. We compare the performance of NPD and NPG under these different conditions. Furthermore, we also test the efficacy of a multivariate extension of NPD, the conditioned NPD, as a means of testing for the effects of common drive and its ability to discriminate between parallel signal routing. Finally, we bring the presented methods to the analysis of empirically recorded data from patients with Parkinson’s disease. Using an example recording, we examine how artificially imposed changes in the signals’ SNR and linear mixing can change the estimate of dFC made between signals recorded from the human cortex and basal-ganglia.

## 2 Methods

### 2.1 Approach

In this study we utilize spectral coherence for estimates of undirected FC, and NPD/NPG for estimates of dFC. We set up models of continuous neural signals with known connectivity architectures parameterized in MVAR coefficients. Confounds such as signal-to-noise and instantaneous mixing are then introduced following simulation of the MVAR process using an observation model. Using coherence, we first establish the existence of coherent frequencies within the modelled data sets. Patterns of connectivity in the models are then recovered using the two dFC metrics (NPD and NPG). As connectivity in the models is known (by design) we analyse how the metrics perform at recovering an accurate estimation of connectivity profiles. Finally, we look at the methods’ application to empirical data when used to estimate the directed functional connectivity between the STN and motor cortex in recordings made from a patient with Parkinson’s disease (PD).

### 2.2 Software for Analysis, Simulations, and Statistics

Data was analysed using a set of custom scripts written in MATLAB R2017a (The Mathworks, Nantucket, MA, USA). Non-parametric directionality was implemented using the Neurospec toolbox (http://www.neurospec.org/). MVAR models were implemented using the BSMART toolbox (Cui et al. 2008) implemented in FieldTrip (Oostenveld et al. 2011b). All scripts for the analyses presented here can be found in a GitHub repository (https://github.com/twestWTCN/NPD_Validate). A full list of script dependencies, toolboxes used, their authors, and links to their original source code used can be found in Appendix I.

### 2.3 Functional Connectivity

#### 2.3.1 Spectra and Coherence

Spectral estimates were made using multi-windowed Fourier estimation utilizing discrete prolate spheroidal sequences as orthogonal windows (Thomson 1982). Data were divided into 1 second segments (n = 250), and estimates were made using taper numbers equal to a smoothing window of ±2 Hz. We computed the magnitude-squared coherence via:

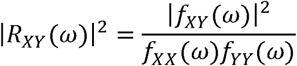

where *f*_*XX*_, *f*_*YY*_, *f*_*XY*_ are the *X* and *Y* autospectra and *XY* cross-spectrum respectively.

#### 2.3.2 Non-Parametric Directionality

Non-parametric directionality provides a model-free estimate of directional correlations within a system through the decomposition of the coherence into components separated by their lags yielding separate instantaneous, forward-lagging, and reverse-lagging spectra (Halliday 2015). This is achieved using pre-whitening of the Fourier transforms. This acts to bring the spectral content of a signal closer to that of white noise, in this case using optimal pre-whitening with minimum mean squared error to compute the whitening filter. This procedure is equivalent to generating two new random processes which have spectra equal to 1 at all frequencies:

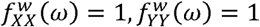

The prewhitening step effectively eliminates the autocorrelation structure of the respective signals but retains bivariate correlations between them. The pre-whitening brings the denominator of the coherence, the product of the autospectra (a normalization factor) equal to 1. Thus, the coherence can be reduced to the cross spectra:

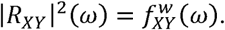

As the coherence loses all terms in the denominator, the equivalent cross-spectrum can then be transformed to the time domain to yield the time-domain correlation function:

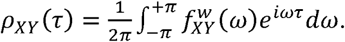

This correlation allows 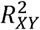 to be decomposed (in the time domain) via Parseval’s theorem by any desired lag. We choose to separate into reverse, instantaneous, and forward components:

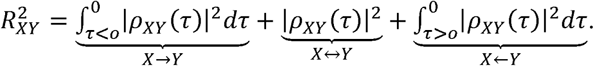

These components may be abbreviated to:

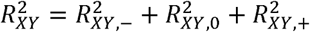

where component 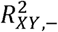 yields correlations in which *Y* lags *X*, 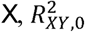 instantaneous correlations, and 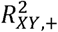 correlations in which *X* lags *Y*. Returning each component back to the frequency domain we obtain 3 measures that sum to yield the original symmetrical coherence:

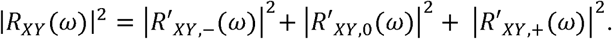

Thus, from each component we can assess spectrally resolved directional interaction whilst accounting for the signals’ autocorrelation structure. For a full derivation of the NPD method and details of its algorithmic implementation please refer to Halliday et al. (2016).

#### 2.3.3 A Multivariate Extension – Conditioned Non-Parametric Directionality

In addition to bivariate NPD, we used a multivariate extension that allows the directional components of coherence to be conditioned upon a third signal (Halliday et al. 2016). The conditionalization of NPD is achieved through a partial regression of *X* and *Y* conditioned on *Z*. This analysis decomposes the partial coherence into the same three directional components: forward, reverse, and zero-lag. This analysis can indicate if information in the bivariate interaction shares variance common to signals in other parts of the network. For example, the partial correlation between *X* and *Y* with *Z* as predictor can be used to determine if the flow of information from *X* → *Y* is independent of area *Z*, or whether the flow of information is *X* → *Z* → *Y*, in which case the partial coherence between *X* and *Y* with *Z* as predictor should be zero. The partial coherence can also be used to investigate if the flow of information is *Z* → *X* and *Z* → *Y*, or if it is *X* → *Y* → *Z* or *Z* → *X* → *Y*, or in the case of common input *Z* → *X and Y*, in which cases the partial coherence, and any directional components, should be zero.

The relationship between the squared coherence function | *R*_*XY*_(*ω*) |^2^ and the squared correlation coefficient was the starting point for the derivation of the non-parametric directionality method in Halliday (2015). The correlation coefficient is given by:

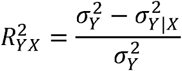

where the conditioned variance, 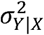 is the variance of the error process following a linear regression of Y on X. It then follows that the correlation coefficient may be conditioned to account for any common effect that a process Z may have on both X and Y by also estimating the residuals following regression with Z:

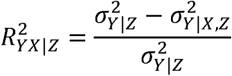

in which the processes X and Y are both conditioned (regressed) against the third process Z. Partial regression is often useful in situations in which it is believed that the tertiary signal Z can account for some or all of the original association between X and Y. Thus, the objective is to distinguish whether there is a genuine correlation 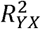 that is distinct from the apparent one induced by 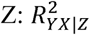. In the same manner by which the correlation coefficient may be conditioned to account for any common effect that a process Z may have on both X and Y, we can condition the estimated coherence | *R*_*XY*_(*ω*) |^2^ on Z:

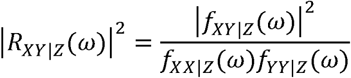

In this way we can form a so called ‘partial’ coherence to determine the association of the coherence between X and Y with predictor Z. By using this form of the coherence as the starting step we can continue with the same decomposition as was made before for bivariate NPD, in order to attain an estimate of the NPD between X and Y but conditioned on Z. In practice we achieve conditioning of the respective autospectra *fXX*|*Z* (*ω*) and *fYY*|*Z* (*ω*) using the approach set out in (Brillinger 1988). This method has been used successfully in LFP recordings to recover known anatomical pathways in the basal-ganglia (West et al. 2018). For full details of the derivation and implementation of conditioned NPD, see Halliday et al. (2016).

#### 2.3.4 Non-Parametric Granger Causality and its Relation to NPD

Granger causality is based on the premise that if a signal *X* causes a signal *Y*, then the past values of *X* can be used to predict the state of *Y* beyond that of the information contained in the past of *Y* alone (Granger 1969). This has conventionally been tested in the context of multivariate autoregressive models fit to the data, and in which the explained variance of *Y* via a ‘restricted’ model based on *Y* alone is compared to that of a ‘full’ model using information of both the past of *X* and *Y* (Geweke 1982). Frequency domain extensions of Granger have been developed (Geweke 1982; Kamiński et al. 2001) and applied widely across many domains of neuroscience (e.g. Brovelli et al. 2004).

The requirement to fit multiple MVAR models can cause several difficulties in analyses, namely: i) the requirement of large model orders to capture complex spectral features; ii) computational cost of model inversion; and iii) assumptions as to the correlation structure of the data in order to capture the signal as an MVAR process. In order to avoid the requirement for the estimation of MVAR models, Dhamala et al. (2008) proposed a non-parametric estimator of Granger Causality. This estimator can be derived from widely used Fourier or wavelet based spectral estimation methods which do not suffer from these complications. The method hinges on the derivation of a spectral matrix directly from the spectral transforms of the data (i.e. Fourier or wavelet) instead of the full transfer and noise covariance matrices specified in an inverted MVAR model. Subsequently, the spectral matrix is factorized to derive the transfer function and noise covariance matrices of the set of signals (Sayed and Kailath 2001). Via this technique it is possible to decompose the total power spectrum of *X* between its intrinsic power and the causal contribution from *Y*. For a full derivation and details of its implementation please refer to Dhamala et al. (2008).

The difference between the way NPG and NPD determine causal or directional components is that NPG uses a decomposition of the signal power into intrinsic and extrinsic components, whereas NPD decomposes a normalised correlation coefficient according to time lag. Both NPG and NPD use a frequency domain approach. The frequency approach in NPG uses the formulation in Geweke (1982) in combination with factorization of the spectral matrix (Wilson, 1972), see Dhamala et al. (1982) for details. NPD is based on the approach of Pierce (1979) to decompose the product moment correlation coefficient and coherence summatively into directional components. The starting point is the spectral matrix (as in NPG). The decomposition is achieved by pre-multiplying the spectral matrix to create an MMSE pre-whitened version, *S*_*w*_ as:

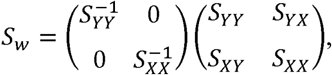

where, *S*_*YY*_ and *S*_*XX*_ are the autospectra, and, *S*_*YX*_ and *S*_*XY*_ are the cross spectra. The effect of this pre-multiplication allows coherence to be calculated directly from the cross-spectra. A normalised correlation coefficient is obtained from the inverse Fourier transform of the MMSE whitened cross spectrum, normalised in the sense that the overall product moment correlation can be obtained through integration of this correlation function. By decomposing the normalised correlation coefficient by time lag and taking the corresponding Fourier transform of each component, NPD creates separate spectra which are used to decompose the coherence summatively (i.e. summing up to the magnitude of the original coherence) for each frequency. The time lags used are forward, reverse and zero-lag decomposing the coherence intro three components. NPD thus decomposes coherence according to time lag in the normalised correlation whereas NPG decomposes the spectrum into intrinsic and extrinsic factors, the presence of non-zero intrinsic factors is taken as indicative of a causal effect in NPG.

#### 2.3.5 Pairwise Versus Multivariate Applications of Metrics

Both NPD and NPG can be used in either a bivariate (pairwise) or a full multivariate (greater than two signals) framework. As pairwise analyses of dFC are by far the most common approach used in the current literature we primarily make a comparison of bivariate NPD and NPG computed between two signals only. However, when investigating issues such as common drive and the influence of tertiary signals we utilize the multivariate NPG (mvNPG) and compare it with conditioned NPD. Conditioned NPD is indicated by the use of brackets to signify the conditioning signal (e.g. NPD(x) signifies NPD conditioned on signal X). This approach is used exclusively in section 3.1 (common drive) and 3.6 (incomplete signals for conditioning).

### 2.4 Simulations

#### 2.4.1 Multivariate Autoregressive (MVAR) Modelling

In order to simulate data that models lagged propagation between simple periodic systems we used multivariate autoregressive (MVAR) modelling. MVAR models are an extension to 1-dimensional autoregressive models in which the present state is dependent upon a linear combination of the previous states plus some stochastic error term. Generally, a P^th^ order MVAR model with .*N* number of states is given by:

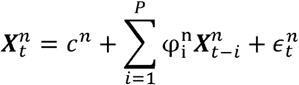

where 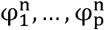 are the autoregressive coefficients for state *n* at lag *i,c^n^* are constants of state *n*, and 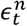 is white noise with zero mean and covariance structure *R*. The AR model takes a model order *P* which determines the number *t* – *P* of previous states of the system that are regressed to form the present state ***X***_*t*_, thus imbuing the system’s evolution with autocorrelation structure with persistence dependent upon *P*. Simple periodic signals may be engineered in the MVAR formulation by setting of alternating signed coefficients at different lags. For example, to obtain a lag two periodicity at state *n* we set 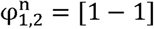. The alternating signs of the coefficients sets up the signal to oscillate with a period equal to the difference in lags. The matrix of coefficients **φ** ∈ ℝ^*N×P*^ specifies the linear terms of a state’s dependency upon its past and neighbouring states. In order to introduce correlations between states we introduced non-zero coefficients off the diagonal. In this way we simulate lagged connectivity by setting positive coefficients between nodes at lags greater than 1. For the parameters of the simulated MVAR models please see appendix II. Simulations were made using the BSMART toolbox. All simulations were run to yield 10^5^ data samples. In order to set a time scale of the simulations we choose an arbitrary sampling frequency of 200 Hz which places simulations around the frequencies typically observed in neural data. The model architecture for each set of figures is outlined using a ball and stick diagram next to the main results.

#### 2.4.2 Observation Modelling

In order to introduce the effects of changes in SNR and instantaneous mixing of signals that can arise due to the practical aspects of experimental recordings of neural signals, we construct an observation model on top of the model of the dynamics:

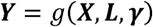

where the observer function *g*()maps from the hidden (dynamical) states ***X*_*t*_** onto the observed states ***Y*_*t*_**. The function utilizes a leadfield matrix ***L*** which performs an instantaneous linear combination of states:

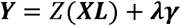

where there is a constraint on the diagonal of L:

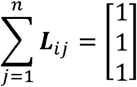

Thus, we can specify the mixing between signals by specifying the off-diagonal entries of matrix ***L***(i.e. ***L***_*i ≠ j*_). When applied to z-normalized, uncorrelated data, the mixing matrix introduces sharedvariance equal to the off-diagonal coefficients of ***L*** (e.g.***L***_12_ = 0.3 is equivalent to 30% shared variance between independent signals 1 and 2).

Furthermore, we model the introduction of noise during signal acquisition by the addition of observation (as opposed to state) noise ***γ*** which we assume to be i.i.d, zero-mean, unit-variance, white noise. Prior to addition of noise the mixed channels are standardised to zero-mean and unit-variance (indicated by function *Z(x)*). Finally, the observer model simulates modulation of the signal-to-noise ratio (SNR) through the gain factor *λ*, where *λ* =0 indicates that there is pure signal. We can then compute the decibel level SNR via:

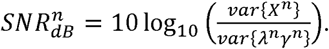

where in the absence of noise 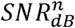 approaches +∞ dB. A 1:1 SNR is equivalent to 0 dB. A large(positive) signal-to-noise (i.e. 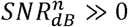) indicates that the data is composed majoritively of signal,whereas a close to zero signal-to-noise (i.e. 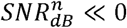) is data comprised predominantly of noise. In some simulations we investigate the role of asymmetric SNR and so report the difference of SNRs between signals:

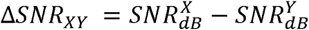

In this case, when Δ*SNR*_*XY*_ = 0, then the signals *X* and *Y* have an equal SNR. When there is an asymmetry in the signals’ gain such that node 2 has the strongest signal (i.e. 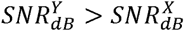) then Δ*SNR*_*XY*_ < 0. Assuming one signal is held constant then a difference in SNRs of 10 dB is equivalent to 10 times increase in the noise in the other signal, 20 dB equivalent to 100 times increase, etc. Because the noise is constructed into the model as additive observation noise, its variance (independent of the simulated signal) is known explicitly. In empirical data, this is a much harder quantity to estimate (Parkkonen 2010).

### 2.5 Experimental Data

#### 2.5.1 Experimental Protocol

In the final experiment of this paper, we investigate how the two dFC metrics (NPD and NPG) perform when estimating the dFC between the cerebral cortex (supplementary motor area; SMA) and the subthalamic nucleus (STN). This connection has been reported to be predominantly cortically leading in patients with Parkinsonism (as estimated with NPG in Litvak et al. 2011, 2012). In this paper we use an example recording in which NPG reveals a clear directed component from SMA → STN. This recording was taken from a cohort of patients with PD who have undergone surgery for deep brain stimulation (DBS). The experimental data contains recordings made using whole head MEG and simultaneous LFP recordings from DBS electrodes implanted into the STN. The recordings were made for approximately 3 minutes with the patient quietly at rest with their eyes open. Experiments investigated the differences in MEG and LFP activity and connectivity when patients were withdrawn from their usual dose of L-DOPA (OFF) versus the L-DOPA treated state (ON). Patients were not undergoing stimulation with DBS at the time of the recording. The two-time series analysed were 183s in duration with MEG from a right SMA virtual sensor recorded simultaneously with an LFP from the right STN in a PD patient in the OFF state at rest. All experiments were conducted in a study approved by the joint ethics committee of the National Hospital of Neurology and Neurosurgery and the University College London Institute of Neurology. The patient gave their written informed consent. For full details of the surgery, implantation, recording, and experimental paradigm please see Litvak et al. 2011.

#### 2.5.2 Pre-processing

The MEG and LFP signals were first down-sampled to 200 Hz. They were then pre-processed using a high-pass filter (passband at 4 Hz, finite impulse response, two-pass). Recordings were truncated 1s at either end to remove border artefacts arising due to movement and equipment initialization. Finally, data were visually inspected to determine the presence of large abnormalities and high amplitude transients. In the case of the example data used here, none were found.

We performed estimation of the empirical SNR of the signals using the equation in section 2.4.2. We estimated the signal amplitude as the variance of the data but bandpass filtered (two-pass finite impulse response; order optimized) within a frequency band in which there was a significant directed coherence. The amplitude of the noise was estimated by taking the total variance of all frequencies outside that of the signal of interest. This was achieved using a band-stop filter (two-pass finite impulse response; order optimized).

Changes to the SNR, asymmetric SNR, and linear mixing of the empirically derived signals were introduced using the same process as listed in section 2.4.2 and assuming that the empirical data ***X*** is arealization of an unknown stochastic dynamical system *f(x)* such as that specified in the MVAR models. This treatment ignores the fact that the data by necessity of empirical recording have already undergone observation with a transform similar in form to *g(x)* but with unknown parameters regarding the lead-field (mixing matrix) and observation noise. Instead we take the empirical recordings as a ground-truth and investigate subsequent changes following artificially induced confounds.

### 2.6 Statistics

#### 2.6.1 Permutation Confidence Intervals

In order to form confidence intervals for the connectivity metrics we make no assumptions as to the form of their distributions but instead form permutation distributions by resampling without replacement. As both the NPG and NPD are based upon a periodogram estimate of the spectral density we reshuffle the order of the individual Fourier transforms 200 times and then compute the statistic (either NPG or NPD). We take the 99% percentile of this distribution as the P = 0.01 confidence limit. Limits are plot in figures as a dashed line with arrows on the side of the axis to indicate their positions.

#### 2.6.2 Least-Squares Regression

In the case of some confounds the response profiles were found to be sigmoidal functions with a maximum response, midpoint *x*_0_; and steepness *κ*. We used a least-squares regression to fit the logistic function. All reported fits exceeded R^2^ > 0.95 and we report the parameters of the curves as summary statistics of the connectivity metrics’ modulation by a confound.

## 3 Results

### 3.1 Organization of the Results

In the following sections: 3.2, 3.3, 3.4, and 3.5 the effects of common drive, degradation of SNR, asymmetric SNR, and instantaneous signal mixing upon estimation of dFC using NPD and NPG are investigated. In figures 1-4 examples of the impact of these individual confounding factors upon the power and connectivity spectra are presented. In order to summarise the effects of the confounds across a much larger range of scales, in figure 5 A-C the effect of SNR, unequal SNR, and signal mixing are visualized as a plot of the relevant statistic of the connectivity (i.e. strength or asymmetry) against a scale of values of the confounding factor. In each of the following sections (3.2-3.5) we inspect first the example spectra displayed in figures 1-4, and then go on to establish the total effect over the full range of the confound using the data illustrated in figure 5 A-C.

**Figure 1.**
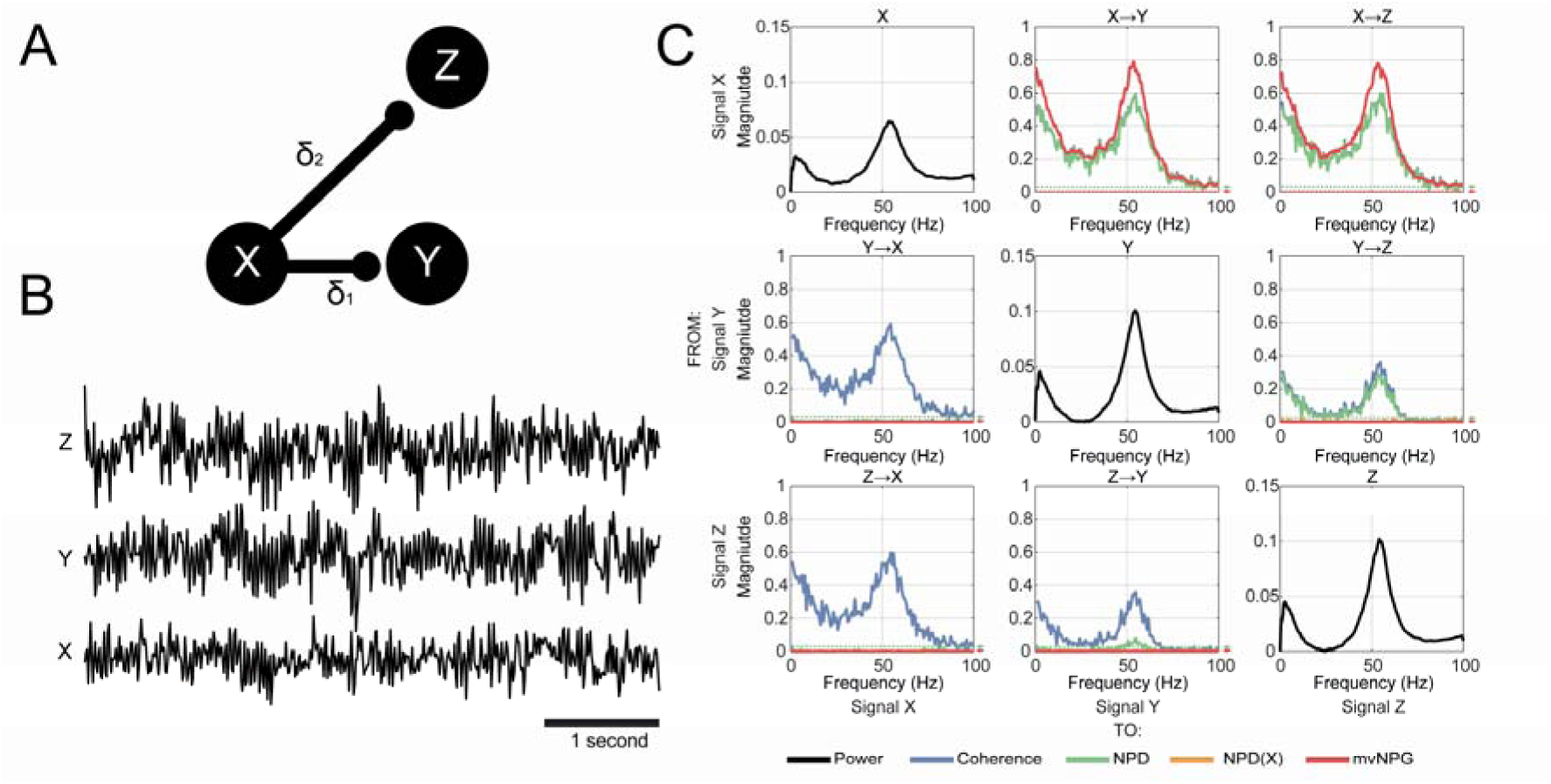
Three -node simulation of MVAR model to compare functional connectivity metrics. **(A)** A simple three state, 3^rd^ order MVAR model was used to simulate coupling of autonomous periodic signals. Connectivity was simulated using non-zero coefficients at lag 2 for node X → Y, and lag 3 for X → Z. Correlations are lagged such that the time delays are unequal (i.e. δ_1_ < δ_2_)**. (B)** Example 5-second realization of the simulated MVAR processes. **(C)** Connectivity matrix of the coupled signals. Autospectra are shown on the diagonal (black). Undirected functional connectivity (coherence) is shown in blue. Estimates of directed connectivity are shown for multivariate nonparametric Granger causality (mvNPG; red); Non-parametric Directionality (NPD; green); and NPD conditioned on signal X (NPD(X); orange). NPD identifies spurious directional connectivity between Y and Z due to the lagged correlations of X → Y relative to X → Z. Spurious connectivity is removed partializing the NPD estimate upon the signal at the common source at node X (NPD(X)) which acts to remove all spurious connectivity. Permutation confidence intervals (P = 0.01) are shown for NPD and mvNPG by the green and red dashed lines and arrows respectively.

### 3.2 Effects of Lagged Dependencies and Common Drive

We first demonstrate the efficacy of the metrics at recovering simple hierarchical architectures and establish how common input can act to confound them. To this end we present results from a simple 3-state, 3^rd^ order MVAR model with no signal mixing and zero observation noise. The MVAR model is imbued with periodic dynamics that are identical at each node and are driven by noise with fixed covariance structure. Non-zero matrix coefficients are all fixed at 0.5 and the full MVAR parameters can be seen in table 1 of appendix II. We design the MVAR model (figure 1A) such that all edges originate at node X and correlations are lagged such that input arriving at node Z lags that at node Y (δ_1_ < δ _2_). This introduces a deliberate confound as dFC methods estimating causality in a way dependent upon temporal lag will assign spurious causality from Y to Z, due to the difference in arrival times of input from X. An example time series of the process is shown in figure 1B and the resulting analyses of the functional connectivity are shown in figure 1C.

This model generates rhythmic activity at ∼60 Hz as indicated by the peaked autospectrum for each node. Functional connectivity as measured using standard coherence shows significant connectivity (> 0.5) between all nodes, albeit reduced for the connection between Y and Z. We next estimate directed connectivity using NPD. NPD shows that all connections are in the forward direction for X → Y and X → Z. As the full coherence is equal to the sum of the directional components, the overlap of the forward NPD (spectra in the upper diagonal of the figure) with the coherence shows that they are equivalent. The shorter lag in transmission from node X → Y compared to X → Z, results in a spurious estimate of coupling from Y → Z when estimated with NPD. However, when we condition the NPD upon the signal that both node X and Z receive common input from (NPD conditioned on node X; NPD(X)), we see that the Y→ Z correlation is abolished.

When pairwise Granger (NPG) was applied to the simulated data, the connectivity from X to Z and Y was very similar in form to the unconditioned NPD (data not shown). However, as the multivariate form of Granger (mvNPG) considers the full covariance across all nodes, its application acted to remove the spurious Y→ Z correlation that arose due to the common drive. This limitation of pairwise NPD is readily overcome using the multivariate extension that allows for the conditioning of the common input. In this way the results between mvNPG and NPD conditioned upon the common input (NPD(X)) are comparable. Both NPD(X) and mvNPG give estimates of Y → Z that are below the P =0.01 confidence interval indicating the absence of any significant directed connectivity between X andY.

### 3.3 Effects of Low Signal-to-Noise Ratios

Recordings of field activity in the brain are made in the presence of both endogenous neural background activity as well as observer noise originating from recording equipment and the external environment. In figure 2 we simulate the effects of signal-to-noise ratio (SNR) upon estimates of functional connectivity. The variance of the MVAR process was clamped and is equal in all simulations. We used additive Gaussian noise in the observation model to simulate an SNR of 1:0 (+ ∞ dB), 4:3 (+ 1.25 dB), and 1:3 (-4.7 dB). All functional connectivity metrics were resistant up to moderate amount of additive noise (SNR_dB_ = + 1.25 dB), but all estimates were heavily attenuated for the greatest noise tested i.e. SNR = -4.7 dB. When looking across a range of SNRs (figure 5A), both NPD and NPG approached 0 when the data became almost entirely noise i.e. SNR approached 0:1 (-∞ dB). Responses were sigmoidal for all three metrics measured with half maximum suppression around 50% signal loss. Non-linear least-squares fitting yielded parameter estimates of the logistic rise for each FC metric (midpoint *x*_0_and steepness *κ*): coherence 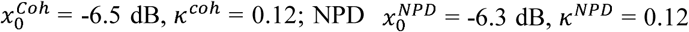 and 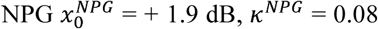. From these estimates and the curves shown in figure 5A, it is clear that coherence and NPD effectively share the same response profile to SNR. NPG is more sensitive to noise with estimates becoming degraded at higher SNRs 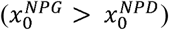. Overall the two metrics have a difference in sensitivities of 8.2 dB, with NPG being more sensitive to noise with almost an order of magnitude greater than NPD. However, the increased variance of the NPD estimator over NPG means that the crossing point of the P = 0.01 confidence intervals occurs at roughly the same SNR for both estimators at around -18 dB. This indicates that either metric will not estimate a statistically significant connection if the signal is comprised of more than 99.5% noise.

**Figure 2.**
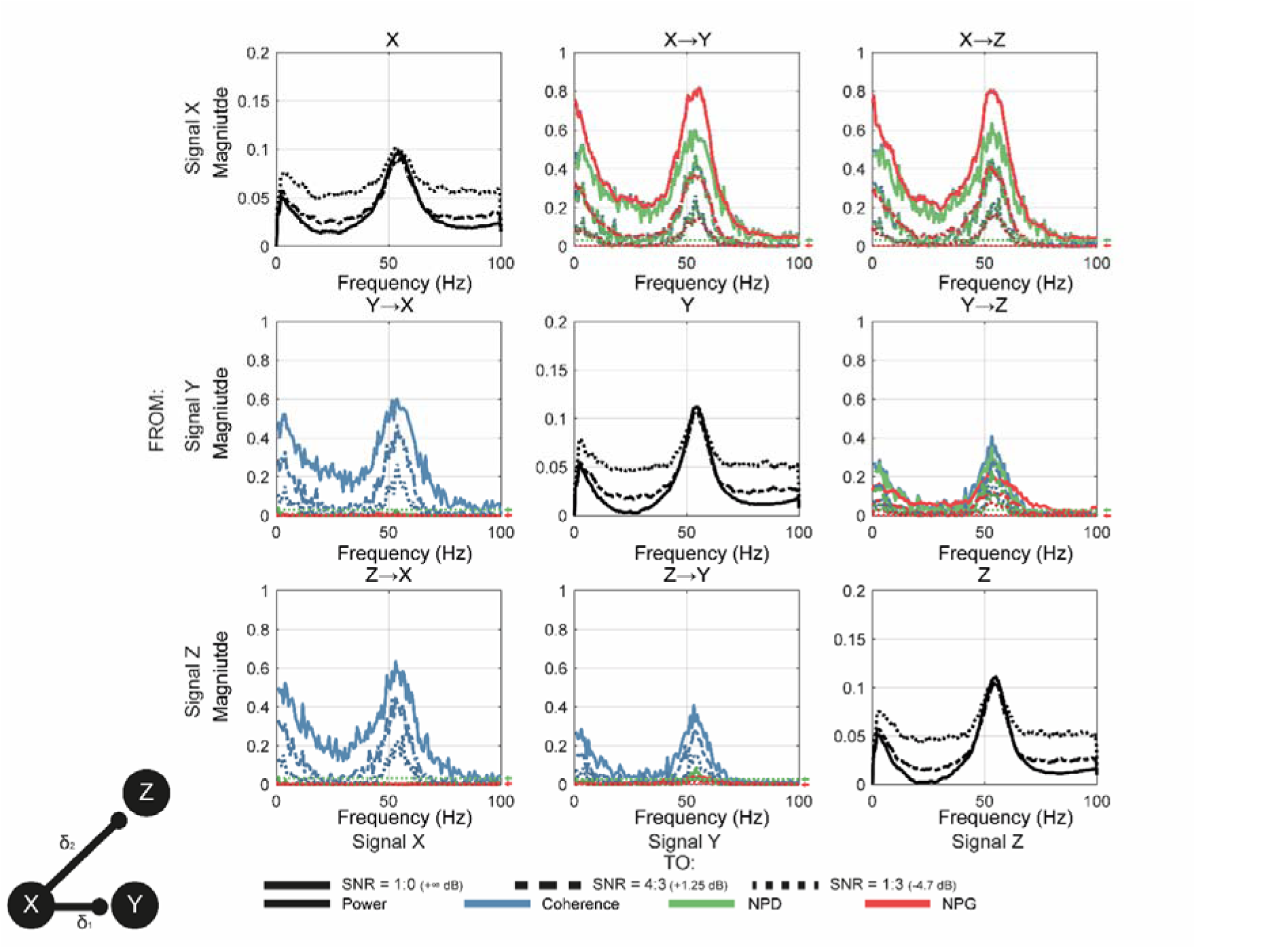
Analysis of the effects of signal-to-noise ratio (SNR) upon estimates of functional connectivity. The confounding effects of poor SNR were simulated by adding Gaussian noise to the MVAR processes and clamping the overall variance equal to 1. The MVAR model is identical in form to that used in figure 1 and its structure is given by the ball and stick diagram in the inset. Simulated SNRs at: 1:0 (+ ∞ dB; bold), 4:3 (+ 1.25 dB; ---), and 1:3 (- 4.7 dB; …). The effects upon coherence (blue), NPD (green), and non-parametric Granger causality (NPG) were investigated. All metrics were effectively reduced by increased levels of noise. Permutation confidence intervals (P = 0.01) are shown for NPD and NPG by the green and red dashed lines and arrows respectively.

### 3.4 Effects of Differences in Signal-to-Noise Ratios between Signals

Asymmetries in the SNR of different signals are known to distort the estimation of dFC using Granger causality (Nolte et al. 2008; Bastos and Schoffelen 2016). We next tested whether this was true for NPD. We simplified the model to contain just 2 nodes that were reciprocally connected with the same lag. Again, the output of the MVAR model was clamped to have unit variance. We then modified the SNR of the first node (X) via the same process as for the previous set of simulations but holding the variance of the noise of the second (Y) node clamped to yield an SNR of +20 dB. Signals were constructed with a difference of SNRs between X and Y (Δ*SNR*_*XY*_) equal to -30 dB, -20 dB, and 0 dB and calculated with respect to the SNR of Y which was held constant. The results of the simulations are shown in figures 3 and 5B. In figure 3 we plot the difference in the estimates of the directed connection (i.e. X → Y minus Y → X; Δ*FC*_*XY*_ < 0) to explore any deviation away from the symmetry designed into the MVAR model (Δ*FC*_*XY*_ ≈ 0).

**Figure 3.**
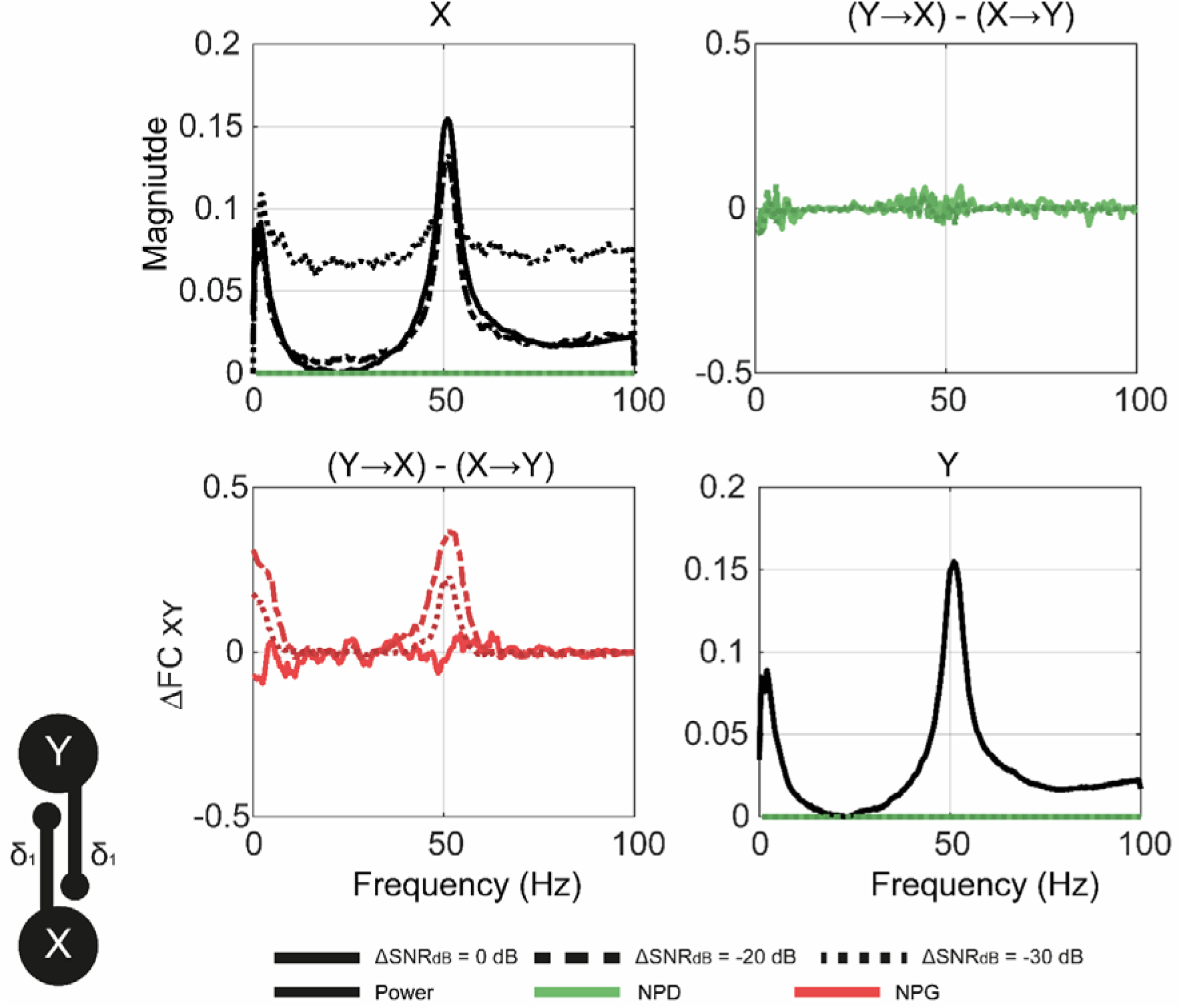
Analysis of the effects of unequal signal-to-noise ratios, measured as a difference of the SNRs between X and Y (Δ*SNR*_*XY*_) upon symmetrical directed functional connectivity (dFC). The confounding effect of connected signals having different SNRs was simulated by the addition of Gaussian noise to signal X but clamping the noise of node Y to + 20 dB. A range of differences in SNR between X and Y (Δ*SNR*_*XY*_) were simulated at 0 dB (X equal SNR to Y; bold), - 20 dB (X 100 times weaker than Y; ---), and - 30 dB (X 1000 times weaker than Y; …). Connectivity was held fixed to be symmetrical. We assessed dFC by plotting the difference in magnitudes of the connectivity for each direction (Δ*FC*_*XY*_) with Δ*FC*_*XY*_≈ 0 as the ground truth. Results from both non-parametric directionality (NPD; green) and non-parametric Granger causality (NPG; red) are shown. In the face of medium amounts of SNR asymmetry, NPG spuriously identifies the strongest signal as the driving node. NPD suffers less from this issue and yields approximately symmetrical estimates for all conditions tested.

Our simulations confirm that NPG is biased by differences in SNR between signals, showing that even at moderate asymmetries (Δ*SNR*_*XY*_ = -20 dB) the weaker signal is estimated to be driven by the stronger i.e. Y→ X. NPD suffers far less from this confound and maintains estimation of the difference in coupling as close to zero for all conditions tested. Analysis with NPD shows far less deviation from the ground truth of symmetrical coupling when the SNRs are unequal. When looking across a range of SNR asymmetries the response of each metric is apparent (figure 5B). NPG spuriously identifies directed coupling, with the bias for Y leading when Δ*SNR*_*XY*_ is in the range -10 dB to -40 dB and peaking around -30 dB (zone II of figure 5C). In contrast, the bias for X leading is when X has a stronger SNR and Δ*SNR*_*XY*_ is in range of +10db to +40 dB, and peaking around Δ*SNR*_*XY*_ = +30 dB (zone IV). At very large (positive) or very small (negative) Δ*SNR*_*XY*_ the bias in coupling is diminished and there is a return to symmetrical estimate of connectivity as both NPD and NPG approach 0 for both directions (zones I and V). However, whilst NPD exhibits a much weaker bias than NPG it does still demonstrate an above significant difference in connectivity. However, deviations in estimation of symmetrical coupling arising due to unequal SNR are roughly an order of difference smaller than NPG with a maximum Δ*NPD*_*XY*_ of -0.045 versus Δ*NPG*_*XY*_ of -0.42.

### 3.5 Effects of Instantaneous Signal Mixing

Neurophysiologically recorded signals such as MEG, EEG, and LFPs are subject to instantaneous mixing of the underlying dipole currents as a result of field spread effects. We next simulate these effects by multiplication of the simulated MVAR process with a linear mixing matrix and investigate the influence of mixing coefficients upon estimates of dFC. We use an identical model to that in section 3.2 (3 state, 3^rd^ order MVAR) but with the addition of the observer model to model signal mixing at a range of values of λ to yield simulations with 0%, 20%, and 60% shared variance. There is no observation noise added. The results of the analysis are shown in figure 4.

**Figure 4.**
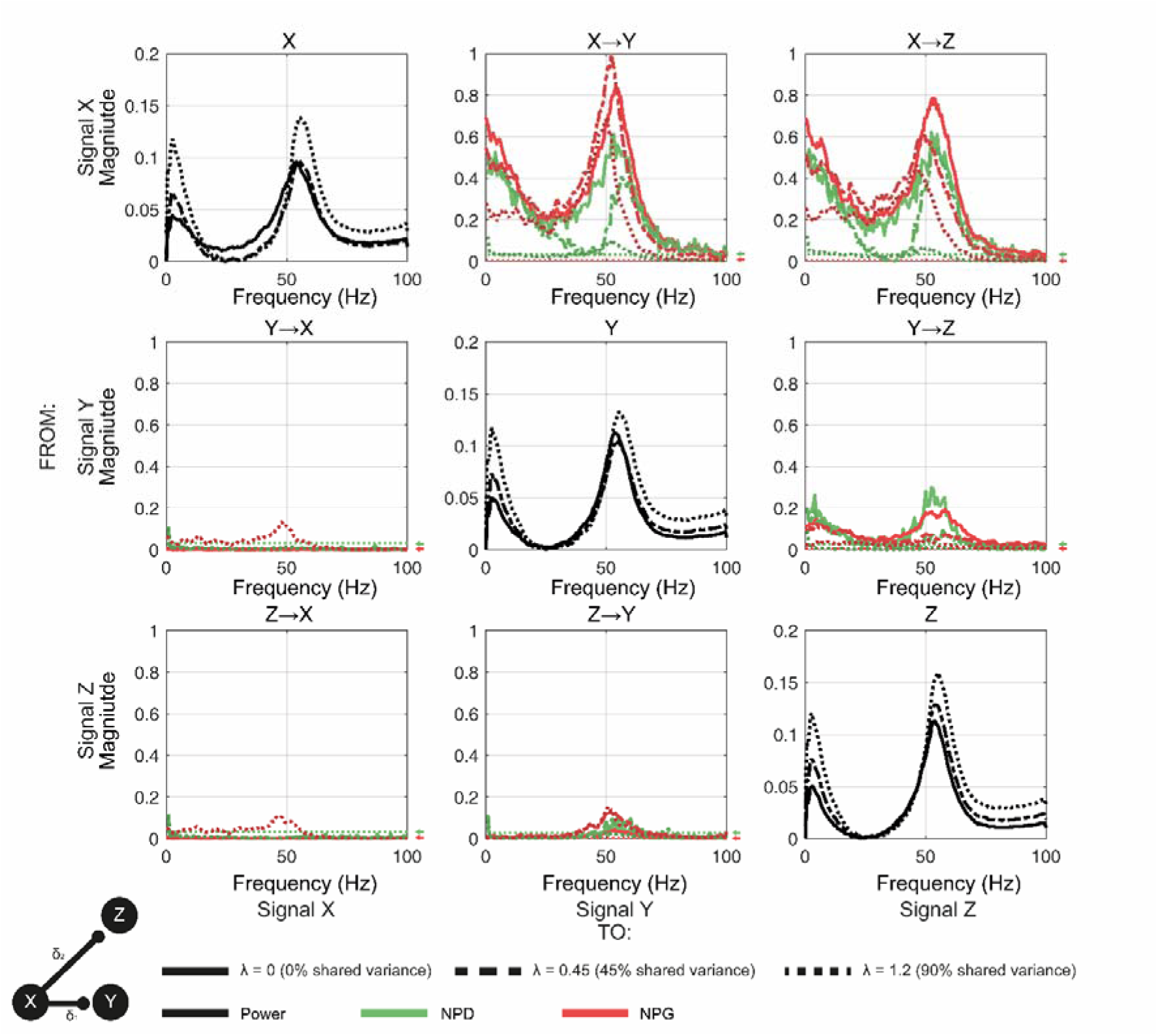
Analysis of the effects of instantaneous mixing upon estimates of directed functional connectivity (dFC). The confounding effects of volume conduction were simulated by multiplication of signals with a mixing matrix with off-diagonal coefficients λ. The unmixed signals were first generated with a three state, 3^rd^ order MVAR model (identical to that used in figures 1 and 2). We simulate three mixing conditions: λ = 0 (zero mixing; bold line), λ = 0.45 (45% shared variance; ---), and λ = 1.2 (90% shared variance; …). dFC is estimated using the lagged components of the NPD (green) or non-parametric Granger (NPG) (red). Permutation confidence intervals (P = 0.01) are shown for NPD and NPG by the green and red dashed lines and arrows respectively.

The confounding effect of instantaneous mixing was established by first estimating the degree to which it may influence the zero-lag component of the NPD. As expected, it was found that the zero-lag NPD is increased by mixing (data not shown), particularly at frequencies outside of the periodic component of the signal. This is by virtue of the fact that the delayed component of the signal is simulated using non-zero, off diagonal model coefficients that result in lagged correlations only at the periodic frequencies. Noise outside these bands introduced by signal mixing readily overcomes the power of the noise independent to each node and leads to zero lag correlations to be greatest at these frequencies.

When using NPD to estimate dFC we found that it accurately reconstructs the designed connectivity up to a moderate degree of signal mixing (45% shared variance), albeit with a reduction of the estimated magnitude of connectivity (e.g. 0.6 to 0.4 for X → Y). At the highest degree of mixing (90% shared variance) the spurious connectivity between nodes Y and Z (introduced by the lagged common drive from node X) becomes increasingly symmetrical with an increase in the connectivity in the reverse direction (i.e. Y → X) despite the absence of these connections in the model arising either by design or by lagged common input. Overall, with increased signal mixing, the estimate of NPD is weakened equally across all connections. Analysis with NPG however shows that mixing has the effect of introducing spurious connectivity between Y and Z, exhibiting a small but significant reversal connectivity at Z → Y at even moderate mixing (45% shared variance). At the greatest degree of mixing, NPG determines statistically significant connections (above P = 0.01 permutation confidence interval) for Y → X and Z → X, neither of which are in the underlying model. Unlike NPD, which shows a uniform reduction in magnitude with increased mixing, the magnitude of NPG estimates depends upon the initial SNR of the nodes. In this instance the X → Z connection is weakened whilst the X → Y is strengthened. This effect is due to the process explored in section 3.4, by which unequal SNR biases the NPG metric.

When testing across a wider range of degrees of signal mixing (figure 5C) the difference in the response of NPD and NPG is apparent. When using NPG to estimate dFC the magnitude of the estimate of X→ Y increases to a maximum at around 65% shared variance and then quickly collapses at very high mixing as instantaneous correlations begin to predominate. This result is related to the findings of the previous section (section 3.4) in which it was shown that signal leakage acts to modify the effective SNR of the signals such that leakage from one signal can act to bias causality to another at moderate amounts of instantaneous mixing. In contrast NPD displays simpler behaviour, as it reduces in amplitude with increased mixing. This occurs because zero-lag coherence predominates, and the lagged components become increasingly small.

**Figure 5.**
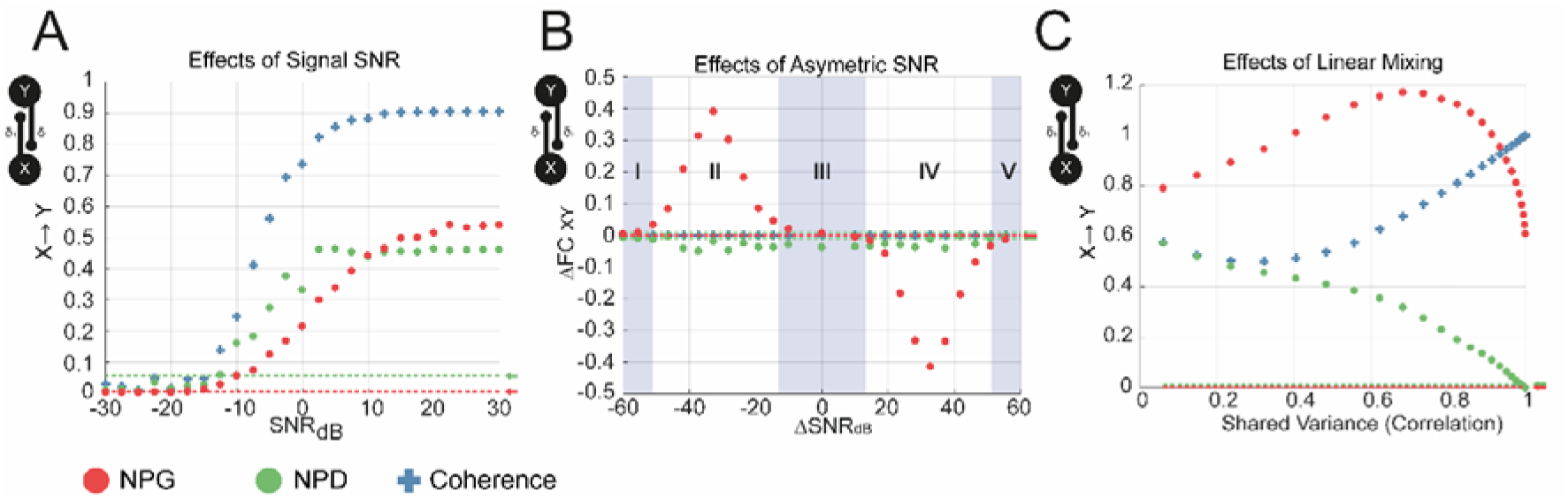
Investigating the effects of signal-to-noise ratios (SNR), SNR asymmetries, and instantaneous linear mixing upon functional connectivity metrics: coherence (blue), nonparametric directionality (NPD; green), and non-parametric Granger causality (NPG; red). Permutation confidence intervals (P = 0.01) are shown for NPD and NPG by the green and red dashed lines and arrows respectively.**(A)** The effect of SNR was tested in the range from -30 dB to +30 dB. All metrics were found to have a sigmoidal response, with half-maximal suppression around SNR = 0 dB. **(B)** The effect of unequal SNR between nodes X and Y (Δ*SNR*_*XY*_) was varied by addition of observation noise to node X or Y separately to yield a range of SNRs from -60 dB to + 60 dB whilst coupling strengths were held fixed. NPG incorrectly identifies asymmetrical coupling for a wide range of Δ*SNR*_*XY*_ (within zone II from -40 dB to -10 dB as well as zone IV from +10 dB to 40 dB). NPD estimates a weak bias towards one signal leading but with differences in directionality remaining close to zero across the range examined. **(C)** The effect of instantaneous signal mixing was examined across a range of mixing coefficients (λ) to yield a range of 0% to 100% shared variance. Coherence is shown to increase as zero-lag correlations predominate with increasing valued λ. The lagged NPD shrinks to zero as instantaneous component of coherence dominates. NPG increases to a maximum at around 65% signal mixing and then sharply falls to zero. Permutation confidence intervals (P = 0.01) are shown for NPD and NPG by the green and red dashed lines and arrows respectively.

### 3.6 Confounds for conditioned directed connectivity arising from incomplete measurement of signals

We next investigate the properties of the multivariate extension to NPD which we term *conditioned* NPD. Conditioned dFC provides a more powerful method with which to explore network functional connectivity; however, in empirical cases, conditioning with a tertiary signal Z may not produce complete attenuation of the spuriously inferred directed connection between X and Y arising from the common input Z. This may arise as a result of: i) incomplete capture of the activity occurring at Z; and/or ii) difference in the routing of signals; and/or iii) because there are other sources of the spuriously inferred connection than Z alone. In cases where structural connectivity is well understood, and the conditioned signal Z is not expected to interconnect the path between nodes X and Y, any attenuation when conditioning can be assumed to arise in information propagated forward in the network (feedforward). On the other hand, if anatomical connectivity is unclear the effect of conditioning upon directed connectivity may also be explained by conventional serial routing (i.e. X→ Z→ Y) but with incompleteness of observed signals at Z resulting in only partial attenuation of the X → Y estimate. In the next set of simulations, we ask whether there are any differences in how the measures of dFC behave in the face of incomplete signal observation.

For this set of simulations, we use a three state, 3^rd^ order MVAR model, with all nodes generating identical autonomous dynamics and identical cross-node coefficients at equal model lags. We test three model connectivities to compare three types of signal propagation: a) serial (i.e. X → Z → Y); b) feedforward (i.e. X → Y → Z); or c) recurrent (i.e. X → Y → Z → X). We simulate incomplete observation of Z by modifying the SNR as in section 3.2. The model architectures and results of simulations are shown in figure 6. We demonstrate that in the simplest case of a serial path, the NPD conditioned on signal Z (NPD(Z)) behaves as expected: the estimate of connectivity X → Y is attenuated as all information between them is routed via Z. With decreasing SNR of the observation of Z we show that conditioning has less and less effect and converges to the estimate yielded by the unconditioned NPD. Pairwise NPD remains constant at all SNRs tested as it does not account for any of the activity at Z. In these simulations, multivariate NPG (mvNPG) was also applied as a way to estimate directed connectivity that accounts for all signals in the model. We find that mvNPG shows a small decrease (∼0.025) in the estimate of X→ Y with increased SNR of Z. This weak attenuation demonstrates that mvNPG can detect serial routing, yet it is not as suited for discriminating directed connectivity (i.e. X→ Y) as when there is relay via a secondary node (X → Z → Y).

**Figure 6.**
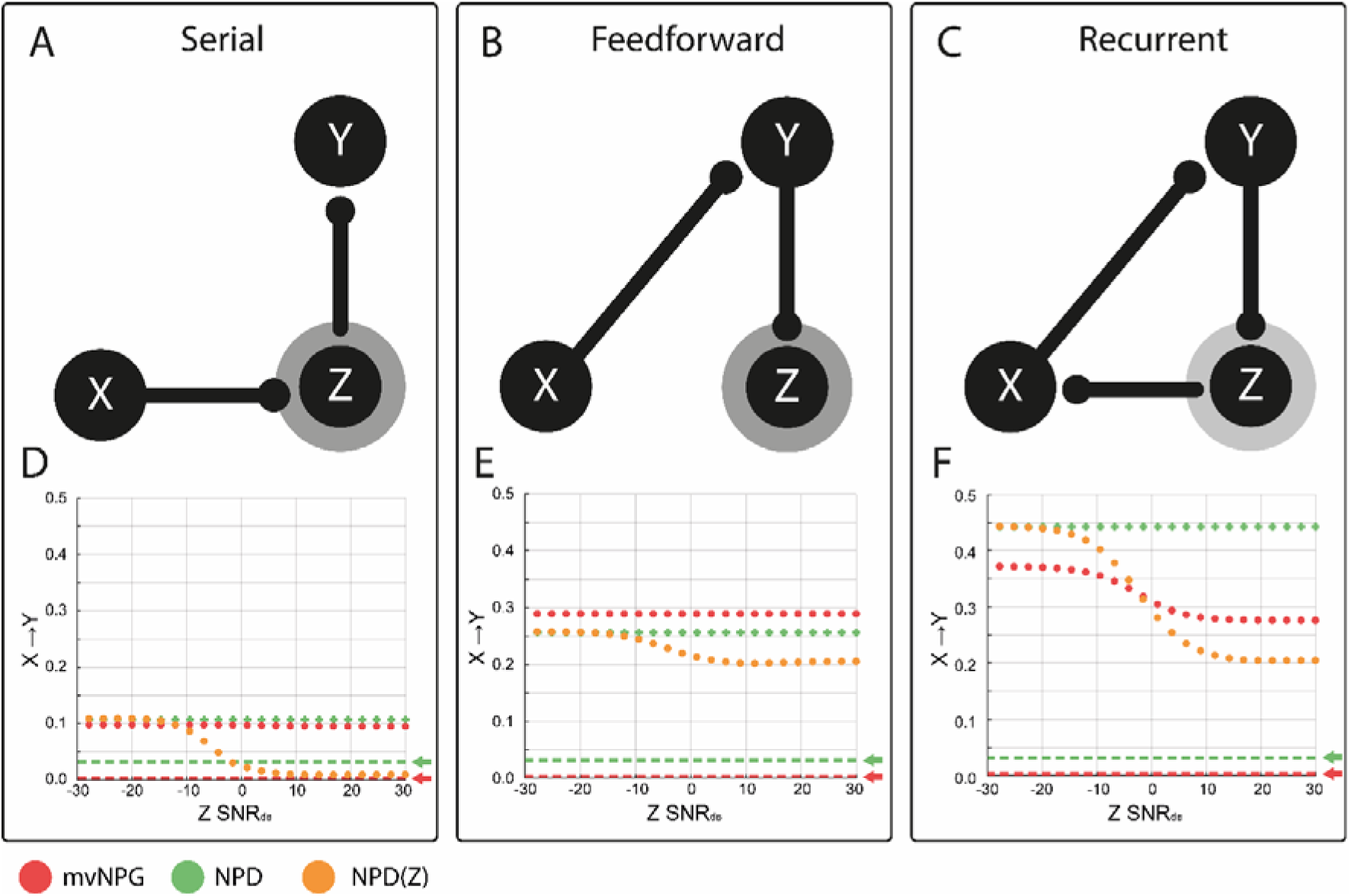
The effects of incomplete signal observation upon estimation of directed functional connectivity: non-parametric Granger causality (NPG); non-parametric directionality (NPD); and NPD conditioned on reference signal Z (NPD(Z)). Simulations investigate the connectivity of X → Y and the influence of propagation involving a tertiary node Z. We simulate incomplete sampling of Z by modifying its signal-to-noise ratio (SNR) via the addition of Gaussian white noise and then clamping the variance equal to 1. **(A and D) Serial propagation –** signals propagate from X→ Z → Y. The results of changing the SNR of Z are shown in panel D. Simulations demonstrate that dFC estimation with NPD/NPG are constant. At complete signal observation (SNR 1:0; +∞ dB), conditioning removes the estimate of dFC. With increasing SNR, the attenuation is diminished to the point where conditioning has no effect. **(B and E) Feed forward connectivity –** signals propagate to feedforward to the tertiary node: X → Y → Z. We find that conditioning has a weak effect (panel E), and the attenuation of NPD(Z) for estimation of X→ Y is again reduced by decreasing SNR of Z. **(C and F) Recurrent connectivity –** a further connection is added to the model to complete a cyclic path in the network: X → Z → Y → X. Decreasing the SNR of Z results in an increased estimation of NPG in X → Y (panel F). We again find that that increased completeness of observation of Z results in an increase in the efficacy of NPD(Z) in determining tertiary (non-direct) signal routing.

We next looked at a feedforward network, where X propagates directly to Y but is then fed on to Z. Because some of the information passed X → Y is contained in Z, we expect conditioning to attenuate the directed connection. Again, we find that NPD(Z) behaves as expected, although the attenuation is weaker than when Z mediated the connection entirely. In this way the difference in values between the NPD and NPD(Z) yields a measure of how much information of X is fed forward from Y→ Z. Thus, decreasing SNR of the observation of Z decreases the attenuating effect of the conditioned NPD. mvNPG remains at a constant magnitude for all SNRs tested. This demonstrates that the multivariate Granger is not sensitive to feedforward configurations whereby the estimation of connectivity between X and Y is not influenced by activity at the terminal (receiving) node.

For the third simulation, we investigated the combination of recurrent loops in the network and incomplete signal observation-two features likely to occur in real recordings from neural systems. We find that with complete signal observation (i.e. SNR →∞) the metrics behave similarly to the feedforward model. A notable difference is the increased NPD of X → Y compared to the feedforward case, as correlations are reinforced by signals resonating across the loop. NPD(Z) behaves in a similar way as before, showing attenuation of the conditioned estimate at low noise levels, but converging back to the unconditioned NPD as the reference signal is obscured by noise and estimation of its confounding influence is lost. The mvNPG estimate of the connection X → Y decreases by 0.1 as the observation noise of Z is reduced. This finding indicates that in the case of recurrent connectivity, mvNPG is sensitive to the quality of the signal recorded at the routing node. In the case of recurrent configurations, this finding shows that mvNPG can readily discriminate between direct X→ Y connectivity and cyclical routing via a secondary signal recorded at Z.

### 3.7 Estimation of Directed Functional Connectivity in Confounded Empirical Data: Cortico-Subthalamic Connectivity

Using the example dataset described in section 2.5 we examine how changes in the overall SNR, differences in SNR between signals, and instantaneous signal mixing may act to confound the estimation of dFC in empirical data recorded from simultaneous MEG and LFP in patients with Parkinson’s disease. We first analyse the original empirical data and then subsequently introduce synthetic confounding effects as described in the methods section that outlined observation modelling. The results of this analysis are presented in figure 7.

**Figure 7.**
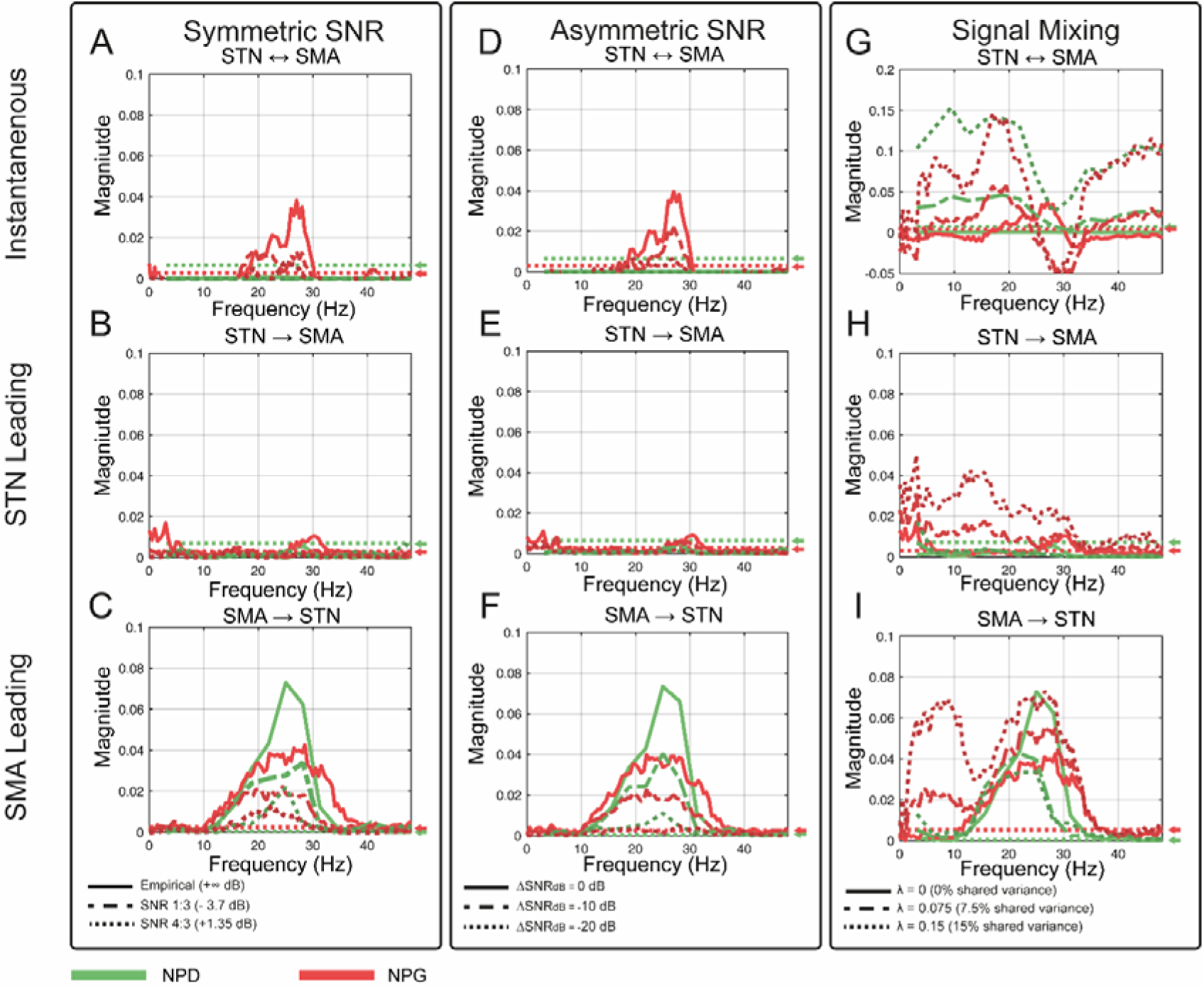
Estimating the confounding effects of symmetric and asymmetric SNR, and instantaneous signal mixing upon experimental data made in patients with Parkinson’s disease. Empirical data is comprised of local field potentials recorded from the STN and a virtual electrode localized to the SMA, computed from whole-head magnetoencephalography. Signals were analysed for dFC using the instantaneous components (first row); STN → SMA (second row); and SMA → STN (third row) parts of the NPD (green) and NPG (red). Empirical data is indicated by bold line; low by the dotted (…); and high degrees by the dashed (---).**(A-C)** The effect of modulating the overall SNR of the signals equally. We used a range of SNRs: 1:1 (+ ∞ dB; bold); 4:3 (+ 1.35 dB; ---); and 1:3 (-4.7 dB; …). **(D-F)** The effect of modulating the SNR of the SMA only. We used a range of Δ*SNR*: 0 dB (bold); -10 dB (---); and -20 dB (…). **(G-I)** The effect of modulating the degree of instantaneous mixing between signals. We simulated a degree of signal mixing: λ = 0 (0% shared variance; bold); λ = 0.075 (7.5% shared variance; ---); and λ = 0.15 (15% shared variance; …).

We demonstrate in the original data that there is a clear asymmetry in coupling with both NPD and NPG indicating a clear dFC from SMA → STN. The zero-lag component (top row figure 7) of the NPD is negligible in the original data. In contrast, the instantaneous component of NPG shows above significance level connectivity at 20-30 Hz. The empirical SNR of the data was estimated using the filtering method described in section 2.5.2. We use activity in the 20-30 Hz band to define the signal (data bandpassed filtered within this range) and then treated all data outside this band as noise (data bandstop filtered at outside the signal frequency range). SNR estimates of the MEG virtual electrode and LFP were -9.5 dB (SNR_SMA_ ≈ 1:9) and -5.0 dB (SNR_STN_ ≈ 1:3) respectively. This yields an empirical Δ*SNR* of -4.5 dB with the LFP measured at the STN having the largest SNR.

In the first set of experiments we reduced the SNR of both signals equally (figure 7 A-C). We added noise to the original signals to yield a range of SNRs: 1:0 (+ ∞ dB); 4:3 (+ 1.35 dB); and 1:3 (-4.7 dB). These analyses show that both NPG and NPD estimates of connectivity respond to a uniform reduction in SNR in a simple and predictable way by reducing their overall magnitude approaching zero as the signals become mostly noise.

Subsequently, the effect of changing the SNR of only one of the signals upon dFC estimates was also investigated (figure 7 D-F). Signals were constructed to have a range of Δ*SNR*: 0 dB; -10 dB; and -20 dB. We reduced the SNR of the SMA signal only 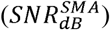, in an attempt to bias the directionality estimates in the reverse direction (i.e. increase the strength of STN → SMA). However, it was found for both NPD and NPG that this had a similar effect to reducing the SNR symmetrically (i.e. when SMA → STN is weakened). This result suggests that for this dataset, it may not be possible to induce a strong bias in the inferred dFC by making one signal weaker than the other (i.e. 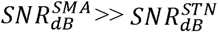) as there is no anatomical STN→SMA feedback (a situation in contrast with simulations investigating asymmetric SNR in section 3.4.).

In the final column of figure 7 (panels G-I) the effect of signal mixing was measured. We simulated a degree of signal mixing: λ = 0 (0% shared variance); λ = 0.075 (7.5% shared variance); and λ = 0.15 (15% shared variance). Again, it was found that the instantaneous component of the NPD behaves as expected, increasing in magnitude with increased signal mixing. This is most apparent in the frequencies outside the main oscillatory bands of activity. When using the instantaneous part of NPG, we found that there was generally an increase, yet the frequencies around the main component (of the peak in the lagged connectivity) were negative and uninterpretable. Furthermore, we show that even moderate increases in the signal mixing (7.5%) corrupt the dFC estimation when using NPG. This is especially apparent at high mixing levels (15%), where a wide band reverse component (STN→SMA) arises, as well as large second peak in the SMA→STN at around 4-12 Hz. NPD estimates are much more stable in comparison and only show a reduction in the original peak with increased mixing, but no spurious peaks emerge outside of this range at any of mixing degrees tested.

## 4 Discussion

The results presented in this paper further support the NPD methodology as an accurate and robust method for the estimation of dFC in continuous neural data. We first provided a face validation of NPD for estimation of the directed interactions between simulated signals. Secondly, we assessed the performance of the NPD metric in the presence of several confounding factors that are likely to arise in experimental recordings of neurophysiological networks, namely: volume conduction, common drive, low SNR, unequal SNRs between signals, and recurrent connectivity. Thirdly, we provided a direct comparison of NPD with a well-established estimate of dFC based on Granger causality – NPG. Finally, our results show that the additional information gained from using a conditioned, multivariate extension of the NPD metric allows for some of the confounding influences of common drive, or nontrivial signal routing, to be mitigated. The degree to which this is achieved is dependent upon the extent to which the signal captures the neural activity at the recording site.

### 4.1 A Summary of Effects of Signal Confounds

#### 4.1.1 Effects of Common Drive

Common input to two parallel neural populations has long been known to be a confounding factor when estimating functional connectivity (Aertsen et al. 1989; Farmer et al. 1993; Horwitz 2003). The limitations of finite sampling over the brain means that no FC metric is immune to this problem as there always remains the potential for an unmeasured common input to the recorded populations from which an FC estimate is made. Our simulations demonstrate this effect where both pairwise NPD and NPG estimates both indicate spurious causality in the case of lagged common input. However, when using multivariate extensions of the two methods, in which the common drive signal is factored out, it is possible to avoid spurious estimation of connectivity between nodes sharing a common drive. This is shown to be true when using the multivariate NPG which accounts for the total covariance across the network. On the other hand, NPD in its simplest form is measured in a pairwise manner and cannot account for the action of a tertiary signal on the naïve estimate. However, we demonstrate that this issue can be remedied using the multivariate extension of NPD in which the influence of a common drive may be regressed out in order to eliminate spurious connectivity between the driven nodes. Whilst this is a solution when the common drive is observed, there still remains the potential confound of an unobserved common signal, to which NPD and NPG are equally susceptible. These issues can be addressed by model based estimators of effective connectivity such as dynamic causal modelling which allow for the inference of unobserved states in a causal network (Friston et al. 2013).

#### 4.1.2 Effects of A/symmetric Signal-to-Noise Ratios

Functional connectivity estimates are subject to the limits of inference implied by the SNR of the available recordings. We demonstrate (figure 5A) that coherence, NPD, and NPG are degraded by poor SNR with similar logistic decays. However, NPG exhibits a greater susceptibility to degradation by noise than NPD. NPG magnitude are reduced at SNRs an order of magnitude higher than those that would elicit an equal reduction in NPD. Despite this, both metrics show a remarkable resistance to even high levels of noise, with the range of SNRs at which both NPG and NPD provide statistically significant estimates of connectivity (i.e. having a magnitude exceeding the P = 0.01 confidence interval) reaching as low as SNR = 1:30 (equivalent to -15 dB). This would suggest that both are robust to the occurrence of false-negative errors as a result of poor SNR in neural recordings. These findings can also explain the common empirical finding of significant functional connectivity in the absence of obvious peaks in the power spectra.

A number of authors have noted that Granger causality is biased by the existence of unequal SNRs (Nolte et al. 2008; Haufe et al. 2013; Bastos and Schoffelen 2016). Our simulations reiterate this fact and demonstrate that NPG is biased to estimate the driving node as the strongest signal (section 3.4; figure 5B). This is an important problem as all neurophysiological signals comprise some unknown mixture of the signal of interest and background noise on a source by source basis. As a result, it can rarely be assumed that the SNRs of two signals are balanced.

This is particularly important when looking at directed connectivity between signals recorded from two different modalities (e.g. MEG and LFP) where the estimate will be biased in favour of the higher gain recording (to lead). For instance, in our data we show a difference in empirical SNRs of 4.5 dB between the LFP and virtual channel signals. This has led some authors to suggest the usage of time-reversed data as surrogate comparison for dFC methods (Haufe et al. 2013) because if a true causal effect is present then time reversal should flip the sign of the directionality. Our simulations demonstrate that estimates made using NPD are less subject to this confound. NPD is still affected by decreased SNR (both asymmetric and symmetric) but shows no bias, as directional estimates decrease uniformly as the SNR goes down. This finding leads us to suggest that in future studies of dFC in multimodal data or in other cases where the signals are likely to be of differing SNR, the NPD method provides a more robust and readily interpretable result over Granger based approaches.

#### 4.1.3 Effects of Simulated Volume Conduction through Signal Mixing

The extent to which signals recorded from the brain are subject to the influence of volume conduction is generally more severe with decreasing distance between the recording electrodes. Experiments have demonstrated that LFPs measured from electrodes separated by a distance of 5 cm will typically show R^2^ values indicating approximately 50% shared variance (Nunez et al. 1997) and so analyses of directed functional connectivity are likely to be significantly affected by instantaneous mixing at distances much closer than this (e.g. recordings made from neighbouring contacts of the same intracranial electrodes). Instead, some authors have shown that functional connectivity analyses are better suited to source localized signals due to the reduced extent of signal leakage (Schoffelen and Gross 2009). This is likely to hold true for the application of NPD analysis to whole brain recordings. It is difficult to find a limit for when zero-lag effects will corrupt a metric such as NPG as this ultimately depends on the nature of the lagged connectivity present in the data. In our simulations, we show that the bias on NPG induced by mixing is dependent upon the original SNR of the signals as mixing introduces some of its confounds by the mechanism of SNR asymmetry discussed in the previous section.

In addition to the benefit of being less susceptible to corruption by volume conduction, NPD provides explicit frequency resolved estimation of the zero-lag component of coherence, making it possible to estimate the extent to which coupling is influenced by instantaneous effects. This characteristic affords NPD an advantage over corrected methods of FC such as imaginary coherence or the phase locking index (Nolte et al. 2004; Vinck et al. 2011) which are set up to ignore zero-phase coherence. In this respect it is important to note that zero-phase coherence can reflect synchronous physiological coupling (Roelfsema et al. 1997).

#### 4.1.4 Effects of Limited Signal Observation

The argument that conditioned metrics of dFC such as conditioned NPD provide an increased ability to infer the causal structure of real-world neural networks hinges upon the assumption that a recorded signal truly captures the complete dynamics of the underlying population through which the signal is routed. In section 3.6 we provide an analysis of how the incomplete observation of signals acts to confound the estimates of dFC under several hypotheses of signal propagation: A) serial; B) feedforward; and C) recurrent connectivity (figure 6). In the case of the simplest architecture - serial propagation, the metrics behave as expected – the less that the signal used to perform the conditioning captures the underlying dynamics, then the less that the conditioning can inform accurate estimation of directed connectivity. In the case of complete signal capture, the conditioning procedure (NPD(Z)) completely attenuates the directed connection (X → Y), as there is no possibility that any of the information contained in Y concerning node X is exclusive of Z. Therefore, if the signal recorded at Z completely captures the dynamics of Y then there is the potential to attenuate the X → Y connection entirely using a conditioning on Z. In the case of feedforward propagation, conditioning will also act to attenuate the estimate, but unlike serial processing (where the reference node Z provides an intermediate node in the chain of propagation between X and Y) the attenuation can never be complete as the variance introduced at Z is not shared with X or Y. In case C we show that the reentrant connection acts to increase the overall coherence due to cyclical passage of information in the circuit. Furthermore, conditioning acts to bring the NPD estimate closer to that of the feedforward model (where the re-entrant connection is missing). In the case of recurrent connectivity, the multivariate NPG also acts to discount the reconnection via Z. When Z is completely captured (SNR is high) then the NPG gives an estimated connectivity equivalent again to the feedforward model.

These observations make it clear that if conditioning removes the inferred connectivity in its entirety then the conditioned node must be in a relay like position (i.e. X → Z → Y). For instance, this was found to be the case in West et al. (2018) where conditioning of NPD between the striatum and subthalamic nucleus upon the external segment of the globus pallidus removed connectivity almost entirely, leading to the conclusion that information propagated serially in the network, a finding in-line with known anatomical details of the indirect pathway of the basal-ganglia. However, the findings described in the present paper regarding the combination of circuit organization and SNR of conditioned signals introduce ambiguity when interpreting the results of conditioned or multivariate estimates of directed connectivity in empirical data. For instance, incomplete attenuation of conditioning may arise either from poor SNR of the reference signal in a serial network or may indicate that the conditioned signal is placed in either a feedforward or recurrent configuration. In this case it is necessary to combine evidence from multiple conditioning steps (e.g. also conditioning X → Z on Y) in order to determine the exact signal routing.

### 4.2 Extensions and Final Conclusions

We have presented a validation of NPD, a novel metric for the assessment of dFC, in continuous neural recordings such as that measured in methods commonly used for human neuroimaging. We argue that in the face of common practical issues arising from the physical limitations of many experimental recording methods, as well as from the complex biology of the systems that they aim to explore, NPD and its conditioned extension provide a useful method that builds upon the founding principles of the more established Granger causality. The NPD measure (conditioned and unconditioned) has been recently demonstrated to provide insights into the patterns of propagating neural activity in animal electrophysiology (West et al. 2018) and is likely to have wide application across other domains of clinical and experimental neuroscience. The finding that NPD is robust to the confounding effects of SNR asymmetry means that it may be readily applied to multi-modal neural recordings without some of the concerns that may arise with Granger-based methods.

The validation provided here is not extensive: there is a wide range of other existing dFC metrics to which we have not made comparison, and so it is possible that other metrics may perform better than NPG (for an extensive comparison of many metrics, not including NPD, see Wang et al. 2014). Granger causality-based methods have become a staple of the dFC toolbox and form the statistical foundation for several methods developed since including the directed transfer function (Kaminski and Blinowska 1991) and partial directed coherence (Baccalá and Sameshima 2001). An adaptation of the directed transfer function aimed at improving estimation of directed connectivity (i.e. *X* → *Y*) introduced by Korzeniewska et al. 2003 may perform better at recovering known patterns of connectivity in the face of common drive than the metrics presented here. However, NPD shows broadly equivalent results to the Granger based measure but exhibits more robust performance in the recovery of complex network topologies in highly confounded data. The NPD metric is inexpensive to compute, makes limited assumptions of the properties of the data, is flexible to the form of the original spectral estimate and is conceptually simple to formulate. It eschews the computationally expensive estimation of model parameters required for estimates such as parametric Granger causality, and directed transfer function, or iterative binning procedures for information-based metrics such as transfer entropy. Overall, NPD provides a simple and compact statistical description of directed dependencies between signals and is readily interpretable, providing the basis for testable hypotheses of causation in real neural systems.

## 5 Acknowledgements

SFF acknowledges salary funding support from the University College London Hospitals Biomedical Research Centre. T.O.W. thanks UCL CoMPLEX for their funding and support for the duration of this project.

## 6 Funding

S.F.F. receives funding from UCLH BRC. Engineering Research Council UK (awards EPSRC EP/F500351/1 to T.O.W). The Wellcome Trust (ref: 204829) through the Centre for Future Health (CFH) at the University of York to D.H. The Wellcome Centre for Human Neuroimaging is supported by core funding from the Wellcome 203147/Z/16/Z to T.O.W and V.L. UK MEG community is supported by the MRC UKMEG Partnership grant MR/K005464/1.

## 7 Competing Interests

The authors have no competing interests to declare.

## 10 Appendices

### 10.1 List of External Toolboxes Used in Analysis, Statistics, and Plotting

1. BSMART toolbox-Hualou Liang, Steven Bressler, Mingzhou Ding (Cui et al. 2008): http://www.brain-smart.org/
2. Fieldtrip toolbox-Donders Insitute, Radboud University (Oostenveld et al. 2011a): www.fieldtriptoolbox.org/
3. Linspecer -David Kun: https://github.com/davidkun/linspecer
4. Neurospec 2.11 toolbox-David Halliday, University of York: http://www.neurospec.org/
5. SPM 12 toolbox-UCL, Wellcome Centre for Human Neuroscience: https://www.fil.ion.ucl.ac.uk/spm/

### 10.2 Table of MVAR Coefficients

| Simulating Common Drive, Signal-to-noise, instantaneous mixing (figure 1, 2, 4, 5C) |  |
| --- | --- |
| Transfer Matrix | Noise Covariance Matrix |
| $X_1 = \begin{bmatrix} 0.5 & 0 & 0 \\ 0 & 0.5 & 0 \\ 0 & 0 & 0.5 \end{bmatrix}$ $X_2 = \begin{bmatrix} -0.5 & 0 & 0 \\ 0.5 & -0.5 & 0 \\ 0 & 0 & -0.5 \end{bmatrix}$ $X_3 = \begin{bmatrix} 0.5 & 0 & 0 \\ 0 & 0.5 & 0 \\ 0.5 & 0 & 0.5 \end{bmatrix}$ | $\epsilon = \begin{bmatrix} 0.3 & 0 & 0 \\ 0 & 0.3 & 0 \\ 0 & 0 & 0.3 \end{bmatrix}$ |
| Simulating Asymmetric Signal-to-noise (figure 3, 5A, 5B) |  |
| Transfer Matrix | Noise Covariance Matrix |
| $X_1 = \begin{bmatrix} 0.5 & 0 \\ 0 & 0.5 \end{bmatrix}$ $X_2 = \begin{bmatrix} -0.5 & 0.35 \\ 0.35 & -0.5 \end{bmatrix}$ $X_3 = \begin{bmatrix} 0.5 & 0 \\ 0 & 0.5 \end{bmatrix}$ | $\epsilon = \begin{bmatrix} 0.3 & 0 \\ 0 & 0.3 \end{bmatrix}$ |
| Simulating Incomplete Signals for Conditioning: Serial (figure 6A, D) |  |
| Transfer Matrix | Noise Covariance Matrix |
| $X_1 = \begin{bmatrix} 0.5 & 0 & 0 \\ 0 & 0.5 & 0 \\ 0 & 0 & 0.5 \end{bmatrix}$ | $\epsilon = \begin{bmatrix} 0.3 & 0 & 0 \\ 0 & 0.3 & 0 \\ 0 & 0 & 0.3 \end{bmatrix}$ |
| $X_2 = \begin{bmatrix} -0.5 & 0 & 0 \\ 0 & -0.5 & 0 \\ 0 & 0 & -0.5 \end{bmatrix}$ $X_3 = \begin{bmatrix} 0 & 0 & 0 \\ 0.3 & 0 & 0 \\ 0 & 0.3 & 0 \end{bmatrix}$ | |
| Simulating Incomplete Signals for Conditioning: Feedforward (figure 6B, E) |  |
| Transfer Matrix | Noise Covariance Matrix |
| $X_1 = \begin{bmatrix} 0.5 & 0 & 0 \\ 0 & 0.5 & 0 \\ 0 & 0 & 0.5 \end{bmatrix}$ $X_2 = \begin{bmatrix} -0.5 & 0 & 0 \\ 0 & -0.5 & 0 \\ 0 & 0 & -0.5 \end{bmatrix}$ $X_3 = \begin{bmatrix} 0.5 & 0 & 0 \\ 0 & 0.5 & 0 \\ 0.3 & 0 & 0.5 \end{bmatrix}$ | $\epsilon = \begin{bmatrix} 0.3 & 0 & 0 \\ 0 & 0.3 & 0 \\ 0 & 0 & 0.3 \end{bmatrix}$ |
| Simulating Incomplete Signals for Conditioning: Recurrent (figure 6C, F) |  |
| Transfer Matrix | Noise Covariance Matrix |
| $X_1 = \begin{bmatrix} 0.5 & 0 & 0 \\ 0 & 0.5 & 0 \\ 0 & 0 & 0.5 \end{bmatrix}$ $X_2 = \begin{bmatrix} -0.5 & 0 & 0 \\ 0 & -0.5 & 0 \\ 0 & 0 & -0.5 \end{bmatrix}$ $X_3 = \begin{bmatrix} 0 & 0.3 & 0 \\ 0 & 0 & 0.3 \\ 0.3 & 0 & 0 \end{bmatrix}$ | $\epsilon = \begin{bmatrix} 0.3 & 0 & 0 \\ 0 & 0.3 & 0 \\ 0 & 0 & 0.3 \end{bmatrix}$ |

## Bibliography

Aertsen AM, Gerstein GL, Habib MK, Palm G. Dynamics of neuronal firing correlation: modulation of “effective connectivity.” J Neurophysiol 61: 900–917, 1989.

Baccalá LA, Sameshima K. Partial directed coherence: a new concept in neural structure determination. Biol Cybern 84: 463–474, 2001.

Bastos AM, Schoffelen J-M. A Tutorial Review of Functional Connectivity Analysis Methods and Their Interpretational Pitfalls. Front Syst Neurosci 9: 175, 2016.

Bressler SL, Seth AK. Wiener–Granger Causality: A well established methodology. Neuroimage 58: 323–329, 2011.

Brillinger DR. Time series: data analysis and theory [Online]. Holt, Rinehart, and Winston. https://books.google.co.uk/books?id=9hzvAAAAMAAJ.

Brillinger DR. Some Statistical Methods for Random Process Data from Seismology and Neurophysiology. Ann Stat 16: 1–54, 1988.

van den Broek S., Reinders F, Donderwinkel M, Peters M. Volume conduction effects in EEG and MEG. Electroencephalogr Clin Neurophysiol 106: 522–534, 1998.

Brovelli A, Ding M, Ledberg A, Chen Y, Nakamura R, Bressler SL. Beta oscillations in a large-scale sensorimotor cortical network: Directional influences revealed by Granger causality. Proc Natl Acad Sci U S A 101: 9849–9854, 2004.

Brunner C, Billinger M, Seeber M, Mullen TR, Makeig S. Volume Conduction Influences Scalp-Based Connectivity Estimates. Front Comput Neurosci 10: 121, 2016.

Cui J, Xu L, Bressler SL, Ding M, Liang H. BSMART: A Matlab/C toolbox for analysis of multichannel neural time series. Neural Networks 21: 1094–1104, 2008.

Dhamala M, Rangarajan G, Ding M. Analyzing information flow in brain networks with nonparametric Granger causality [Online]. Neuroimage 41: 354–362, 2008. https://www.sciencedirect.com/science/article/pii/S1053811908001328 [19 Sep. 2016].

Ding M, Chen Y, Bressler SL. Granger Causality: Basic Theory and Application to Neuroscience. In: Handbook of Time Series Analysis. Wiley-VCH Verlag GmbH & Co. KGaA, p. 437–460.

Farmer SF, Bremner FD, Halliday DM, Rosenberg JR, Stephens JA. The frequency content of common synaptic inputs to motoneurones studied during voluntary isometric contraction in man. J Physiol 470: 127–55, 1993.

Friston K, Moran R, Seth AK. Analysing connectivity with Granger causality and dynamic causal modelling. Curr Opin Neurobiol 23: 172–178, 2013.

Friston KJ. Functional and Effective Connectivity: A Review. Brain Connect 1: 13–36, 2011.

Geweke J. Measurement of Linear Dependence and Feedback Between Multiple Time Series. J Am Stat Assoc 77: 304, 1982.

Granger CWJ. Investigating Causal Relations by Econometric Models and Cross-spectral Methods. Econometrica 37: 424, 1969.

Halliday D, Rosenberg JR, Amjad A, Breeze P, Conway BA, Farmer SF. A framework for the analysis of mixed time series/point process data—Theory and application to the study of physiological tremor, single motor unit discharges and electromyograms. Prog Biophys Mol Biol 64: 237–278, 1995.

Halliday DM. Nonparametric directionality measures for time series and point process data. J Integr Neurosci 14: 253–277, 2015.

Halliday DM, Senik MH, Stevenson CW, Mason R. Non-parametric directionality analysis – Extension for removal of a single common predictor and application to time series. J Neurosci Methods 268: 87–97, 2016.

Haufe S, Nikulin V V., Müller K-R, Nolte G. A critical assessment of connectivity measures for EEG data: A simulation study. Neuroimage 64: 120–133, 2013.

Haufe S, Nikulin V V, Nolte G. Alleviating the Influence of Weak Data Asymmetries on Granger-Causal Analyses. In: Latent Variable Analysis and Signal Separation, edited by Theis F, Cichocki A, Yeredor A, Zibulevsky M. Berlin, Heidelberg: Springer Berlin Heidelberg, 2012, p. 25–33.

Horwitz B. The elusive concept of brain connectivity. Neuroimage 19: 466–470, 2003.

Kajikawa Y, Schroeder CE. How local is the local field potential? Neuron 72: 847–58, 2011.

Kaminski M, Blinowska KJ. The Influence of Volume Conduction on DTF Estimate and the Problem of Its Mitigation. Front Comput Neurosci 11: 36, 2017.

Kamiński M, Ding M, Truccolo WA, Bressler SL. Evaluating causal relations in neural systems: Granger causality, directed transfer function and statistical assessment of significance. Biol Cybern 85: 145–157, 2001.

Kaminski MJ, Blinowska KJ. A new method of the description of the information flow in the brain structures. Biol Cybern 65: 203–210, 1991.

Korzeniewska A, Mańczak M, Kamiński M, Blinowska KJ, Kasicki S. Determination of information flow direction among brain structures by a modified directed transfer function (dDTF) method. J Neurosci Methods 125: 195–207, 2003.

Litvak V, Eusebio A, Jha A, Oostenveld R, Barnes G, Foltynie T, Limousin P, Zrinzo L, Hariz MI, Friston K, Brown P. Movement-related changes in local and long-range synchronization in Parkinson’s disease revealed by simultaneous magnetoencephalography and intracranial recordings. J Neurosci 32: 10541–53, 2012.

Litvak V, Jha A, Eusebio A, Oostenveld R, Foltynie T, Limousin P, Zrinzo L, Hariz MI, Friston K, Brown P. Resting oscillatory cortico-subthalamic connectivity in patients with Parkinson’s disease. Brain 134: 359–374, 2011.

Nalatore H, Ding M, Rangarajan G. Mitigating the effects of measurement noise on Granger causality. Phys Rev E 75: 31123, 2007.

Newbold P. Feedback Induced by Measurement Errors. Int Econ Rev (Philadelphia) 19: 787, 1978.

Nolte G, Bai O, Wheaton L, Mari Z, Vorbach S, Hallett M. Identifying true brain interaction from EEG data using the imaginary part of coherency. Clin Neurophysiol 115: 2292–307, 2004.

Nolte G, Ziehe A, Nikulin V V., Schlögl A, Krämer N, Brismar T, Müller K-R. Robustly Estimating the Flow Direction of Information in Complex Physical Systems. Phys Rev Lett 100: 234101, 2008.

Nunez PL, Srinivasan R, Westdorp AF, Wijesinghe RS, Tucker DM, Silberstein RB, Cadusch PJ. EEG coherency: I: statistics, reference electrode, volume conduction, Laplacians, cortical imaging, and interpretation at multiple scales. Electroencephalogr Clin Neurophysiol 103: 499–515, 1997.

Oostenveld R, Fries P, Maris E, Schoffelen J-M, Oostenveld R, Fries P, Maris E, Schoffelen J-M. FieldTrip: Open Source Software for Advanced Analysis of MEG, EEG, and Invasive Electrophysiological Data. Comput Intell Neurosci 2011: 1–9, 2011a.

Oostenveld R, Fries P, Maris E, Schoffelen JM. FieldTrip: Open source software for advanced analysis of MEG, EEG, and invasive electrophysiological data. Comput Intell Neurosci 2011, 2011b.

Parkkonen L. Instrumentation and data preprocessing. MEG An Introd. to methods..

Pierce DA. Signal Extraction Error in Nonstationary Time Series. Ann Stat 7: 1303–1320, 1979.

Richter CG, Coppola R, Bressler SL. Top-down beta oscillatory signaling conveys behavioral context in early visual cortex. Sci Rep 8: 6991, 2018.

Roelfsema PR, Engel AK, König P, Singer W. Visuomotor integration is associated with zero timelag synchronization among cortical areas. Nature 385: 157–161, 1997.

Sayed AH, Kailath T. A survey of spectral factorization methods. Numer Linear Algebr with Appl 8: 467–496, 2001.

Schoffelen J-M, Gross J. Source connectivity analysis with MEG and EEG. Hum Brain Mapp 30: 1857–1865, 2009.

Sporns O. Networks of the Brain [Online]. MIT Press. https://books.google.co.uk/books?id=v1DBKE7-UrYC.

Srinivasan R, Winter WR, Ding J, Nunez PL. EEG and MEG coherence: Measures of functional connectivity at distinct spatial scales of neocortical dynamics. J Neurosci Methods 166: 41–52, 2007.

Stam CJ, Nolte G, Daffertshofer A. Phase lag index: Assessment of functional connectivity from multi channel EEG and MEG with diminished bias from common sources. Hum Brain Mapp 28: 1178–1193, 2007.

Van de Steen F, Faes L, Karahan E, Songsiri J, Valdes-Sosa PA, Marinazzo D. Critical Comments on EEG Sensor Space Dynamical Connectivity Analysis. Brain Topogr. (November 30, 2016). doi:10.1007/s10548-016-0538-7.

Swanson LW. Brain Architecture: Understanding the Basic Plan [Online]. OUP USA. https://books.google.co.uk/books?id=4rAlwYWVEroC.

Thomson DJ. Spectrum estimation and harmonic analysis. Proc IEEE 70: 1055–1096, 1982.

Truccolo WA, Ding M, Knuth KH, Nakamura R, Bressler SL. Trial-to-trial variability of cortical evoked responses: implications for the analysis of functional connectivity. Clin Neurophysiol 113: 206–226, 2002.

Vinck M, Oostenveld R, van Wingerden M, Battaglia F, Pennartz CMA. An improved index of phase-synchronization for electrophysiological data in the presence of volume-conduction, noise and sample-size bias. Neuroimage 55: 1548–1565, 2011.

Wang HE, Bénar CG, Quilichini PP, Friston KJ, Jirsa VK, Bernard C. A systematic framework for functional connectivity measures. Front Neurosci 8: 405, 2014.

West TO, Berthouze L, Halliday DM, Litvak V, Sharott A, Magill PJ, Farmer SF. Propagation of Beta/Gamma Rhythms in the Cortico-Basal Ganglia Circuits of the Parkinsonian Rat. J. Neurophysiol. (January 10, 2018). doi:10.1152/jn.00629.2017.

Wiener N. Nonlinear Prediction and Dynamics [Online]. In: Proceedings of the Third Berkeley Symposium on Mathematical Statistics and Probability, Volume 3: Contributions to Astronomy and Physics University of California Press, p. 247–252. https://projecteuclid.org/euclid.bsmsp/1200502197.

